# DNA2 and FANCM function in two distinctive pathways in disrupting TERRA R-loops and suppressing replication stress at ALT telomeres

**DOI:** 10.1101/2025.05.22.655602

**Authors:** Ashwin Ragupathi, Heba Z. Abid, Yun Chen, Rongwei Zhao, Settapong T Kosiyatrakul, Derin I. Yetil, Julia Neiswander, Rachel Feltman, Shannon Thomas, Benjamin Yusupov, Manrose Singh, Li Zheng, Binghui Shen, Huaiying Zhang, Hsueh-Ping C. Chu, Carl L. Schildkraut, Ming Xiao, Dong Zhang

## Abstract

Cancers maintain their telomeres through two main telomere maintenance mechanisms (TMMs): 85-90% of cancers rely on telomerase, while 10-15% of cancers adopt the Alternative Lengthening of Telomere (ALT) pathway. Previously, we and others reported that FANCM, one of the Fanconi Anemia proteins, plays a critical role in suppressing replication stress and DNA damage at ALT telomeres by actively disrupting TERRA R-loops [1–4]. Here, we showed that inactivation of DNA2 in ALT-positive (ALT+) cells, but not in telomerase-positive (TEL+) cells, induces a robust increase of replication stress and DNA damage at telomeres, which leads to a pronounced increase of many ALT properties, including telomere dysfunction-induced foci (TIFs), ALT-associated PML bodies (APBs), and C-circles. We further demonstrated that depletion of DNA2 induces a pronounced increase of TERRA R-loops and a decrease in replication efficiency at ALT telomeres. Most importantly, we uncovered a strong additive genetic interaction between DNA2 and FANCM in the ALT pathway. Furthermore, co-depletion of DNA2 and FANCM causes synthetic lethality in ALT+ cells, but not in TEL+ cells, suggesting that targeting DNA2 and FANCM could be a viable strategy to treat ALT+ cancers. Finally, utilizing the single-molecule telomere assay via optical mapping (SMTA-OM) technology, we thoroughly characterized genome-wide changes in DNA2 deficient cells and FANCM deficient cells and found that most chromosome arms manifested increased telomere length. Unexpectedly, we uncovered many chromosome arm-specific telomere changes in those cells, suggesting that telomeres at different chromosome arms may regulate and respond to replication stress differently. Collectively, our study not only shed new light on the molecular mechanisms of the ALT pathway, but also discovered a new strategy for targeting ALT+ cancer.

## Introduction

Faithful replication and repair of its genome are vital for the fitness and health of a mammalian cell. Throughout the genome, replisomes frequently encounter various impediments, especially at the difficult-to-replicate (DTR) regions, such as telomeres, centromeres, rDNA loci, and common fragile sites [5–7]. Transient pausing, stalling, or even collapsing of replication forks trigger a series of signaling transduction events, known as replication stress response [8, 9]. The known endogenous DNA replication impediments include: (1) unrepaired DNA lesions; (2) mis-incorporated ribonucleotides; (3) repetitive DNA sequences that are prone to form various secondary structures (e.g., G-quadruplex or G4); (4) a collision of a replication fork with the transcription machinery (i.e., transcription-replication conflicts, or TRC); (5) DNA-protein complexes; (6) tightly packed genomic regions (e.g., heterochromatin); (7) R-loops that are formed when a single-stranded RNA displaces one strand of double-stranded DNA (dsDNA) and pairs with the other one. Due to the prevalence of one or more of these DNA replication impediments at the DTRs, replisomes are more prone to pause and stall at the DTRs [5–7]. To maintain a stable genome, mammalian cells have evolved a variety of strategies to deal with replication stress and ensure successful DNA replication in each cell cycle [8–11].

In cancers, telomeres are maintained by two telomere maintenance mechanisms (TMMs): (1) 85-90% of cancers reactivate the telomerase (TEL); and (2) 10-15% of cancers rely on the alternative lengthening (ALT) pathway [12–15]. The unique sequence and structural features of mammalian telomeres pose constant challenges to replisomes. First, one of the strands of telomere is rich in guanines (i.e., the G-rich strand) and is prone to form G4s [16, 17]. Second, the telomeric long noncoding RNA, TERRA, tends to form R-loops with telomeric DNA, which are more prevalent in ALT+ cells than TEL+ cells [18–20]. Third, Drosopoulos and colleagues showed that though DNA replication can occasionally be initiated within telomeres, it predominantly occurs in the adjacent subtelomeric regions, indicating a general paucity of replication origins within telomeres [21]. Moreover, Sfeir and colleagues showed that treatment with a low dose of the reversible DNA polymerase inhibitor, aphidicolin, dramatically increases incidence of fragile telomeres [22]. Collectively, these observations indicate that telomeres may be a special type of common fragile site [23, 24]. Additionally, ALT telomeres are more heterogeneous in length and some of them are quite long [25–27]. Therefore, ALT telomeres may be even more prone to express fragility. Fourth, heterochromatins are enriched at subtelomeres and telomeres [28]. Finally, to protect the ends of linear chromosomes from being recognized as double-stranded DNA breaks (DSBs), there are a variety of high-order DNA-DNA and DNA-protein structures at telomeres. For example, the telomeric G-rich strand overhang folds back and invades the internal double-stranded regions of telomeres, forming the telomere-loop (T-loop) and the displacement-loop (D-loop) [29, 30]. The T-loop and D-loop are further stabilized by the Shelterin complex [31, 32]. These various high-order DNA-DNA and DNA-protein structures could also slow down replisomes. Taken together, mammalian telomeres pose a greater challenge to replisomes and frequently experience heightened replication stress. Indeed, spontaneous DNA damages are detected at telomeres, especially in ALT+ cells [22, 33, 34].

Although the ALT pathway was discovered nearly thirty years ago [25, 26], the detailed mechanism warrants further investigation. ALT+ cells manifest certain cellular and molecular properties that collectively distinguish them from TEL+ cells [13, 14, 35, 36]. The known ALT properties include: (1) increased telomere dysfunction induced foci (TIFs), defined as colocalization of a DNA damage marker, such as γH2AX or 53BP1, with telomeres; (2) increased ALT-associated PML bodies (APBs), defined as colocalization of PML with telomeres; (3) increased levels of extrachromosomal telomeric repeats (ECTRs), including C-circles; (4) increased heterogeneity in the length of telomeres; (5) increased telomeric sister chromatin exchange (tSCE); and (6) the presence of unique telomeric sequence variants [37]. Many DNA damage response and DNA repair proteins also play important roles in the ALT pathway [13, 36, 38]. For example, we and others reported that FANCM, one of the Fanconi Anemia proteins, suppresses replication stress at ALT telomeres by actively disrupting the formation of TERRA R-loops and promotes the survival of ALT+ cells [1–4]. Depletion of FANCM induced a drastic increase of many ALT properties, indicating that FANCM is an important players in the ALT pathway [1–4].

DNA2 is a highly conserved eukaryotic protein and plays a crucial role in both DNA replication and DNA repair [39, 40]. For instance, DNA2 participates in the removal of the DNA flaps generated during lagging strand DNA synthesis, facilitates the long-range DNA end resection during homology-dependent repair (HDR), and remodels the stalled replication forks to facilitate their recovery. DNA2 has also been implicated in telomere biology. For example, the budding yeast DNA2 associates with telomeres in G1 and G2 [41]. In mammalian cells, DNA2 interacts with TRF1, a component of the Shelterin complex. Removing one copy of DNA2 in mouse embryonic fibroblast cells induces a slight increase of fragile telomeres [42]. Using a unique TEL+ cell line (HeLa LT) with super longer telomeres, O’Sullivan and colleagues demonstrated that depletion of DNA2 in the ASF1-deficient cells further increases the formation of C-circles [43].

Here, we investigated the role of DNA2 in the ALT pathway. We found that like FANCM, DNA2 plays a critical role in suppressing replication stress at ALT telomeres by preventing the accumulation of TERRA R-loops. Most importantly, we discovered a strong additive genetic interaction between DNA2 and FANCM. Furthermore, we demonstrated that co-depletion of DNA2 and FANCM causes synthetic lethality in the ALT+ cells, but not in TEL+ cells, suggesting that DNA2 and FANCM could be a valid therapeutic target for treating ALT+ cancer.

## Results

### Depletion of DNA2 in ALT+ cells induces a robust replication stress response and DNA damage at telomeres and upregulates many ALT properties

In previous studies, we and others identified FANCM as one of the most important players in the ALT pathway and demonstrated that FANCM functions to suppress replication stress at ALT telomeres by actively disrupting TERRA R-loops [1–4]. In searching for new factors in the ALT pathway, we focused on DNA2, a well-established DNA replication and DNA repair protein [39, 40]. We knocked down human DNA2 using three different siRNAs in U2OS, an ALT+ osteosarcoma cell line (Fig 1A). The three DNA2 siRNAs slightly reduced cells in G1, while moderately increased cells in S or G2 (Fig S1). We then monitored replication stress response at telomeres using antibodies that recognize the phosphorylated serine-345 of Chk1 (pChk1) and phosphorylated serine-4 and serine-8 of RPA32 (pRPA). Antibodies recognizing TRF1 or TRF2, two subunits of the Shelterin complex [31], were used as the marker for telomeres. As seen in Figs 1B and 1C, depletion of DNA2 in U2OS induced a pronounced increase of pChk1 foci colocalizing with TRF1, suggesting that DNA2 deficiency induces replication stress at ALT telomeres. Consistent with the siDNA2 results, when we treated U2OS cells with two small molecule inhibitors of DNA2, d16 and C5, both of which interfere with the binding of DNA2 to DNA [44, 45], we observed an increase in the pChk1 foci (Figs S2A, S2B, S3A, and S3B) and pRPA foci (Figs S2C and S2D) that colocalized with TRF2. Moreover, when we depleted DNA2 using the same three siRNA in Saos2 (Fig S4A), another ALT+ osteosarcoma cell line, we observed a robust increase of the pChk1 foci colocalizing with TRF1 (Figs S4B and S4C). In contrast, when we depleted DNA2 using the same three siRNA in A549 and HeLa, two TEL+ cells, we did not observe any pronounced increase of pChk1 foci or pRPA foci at telomeres (data not shown), suggesting that DNA2 deficiency induced replication stress response at telomeres is specific for ALT+ cells.

**Figure 1.**
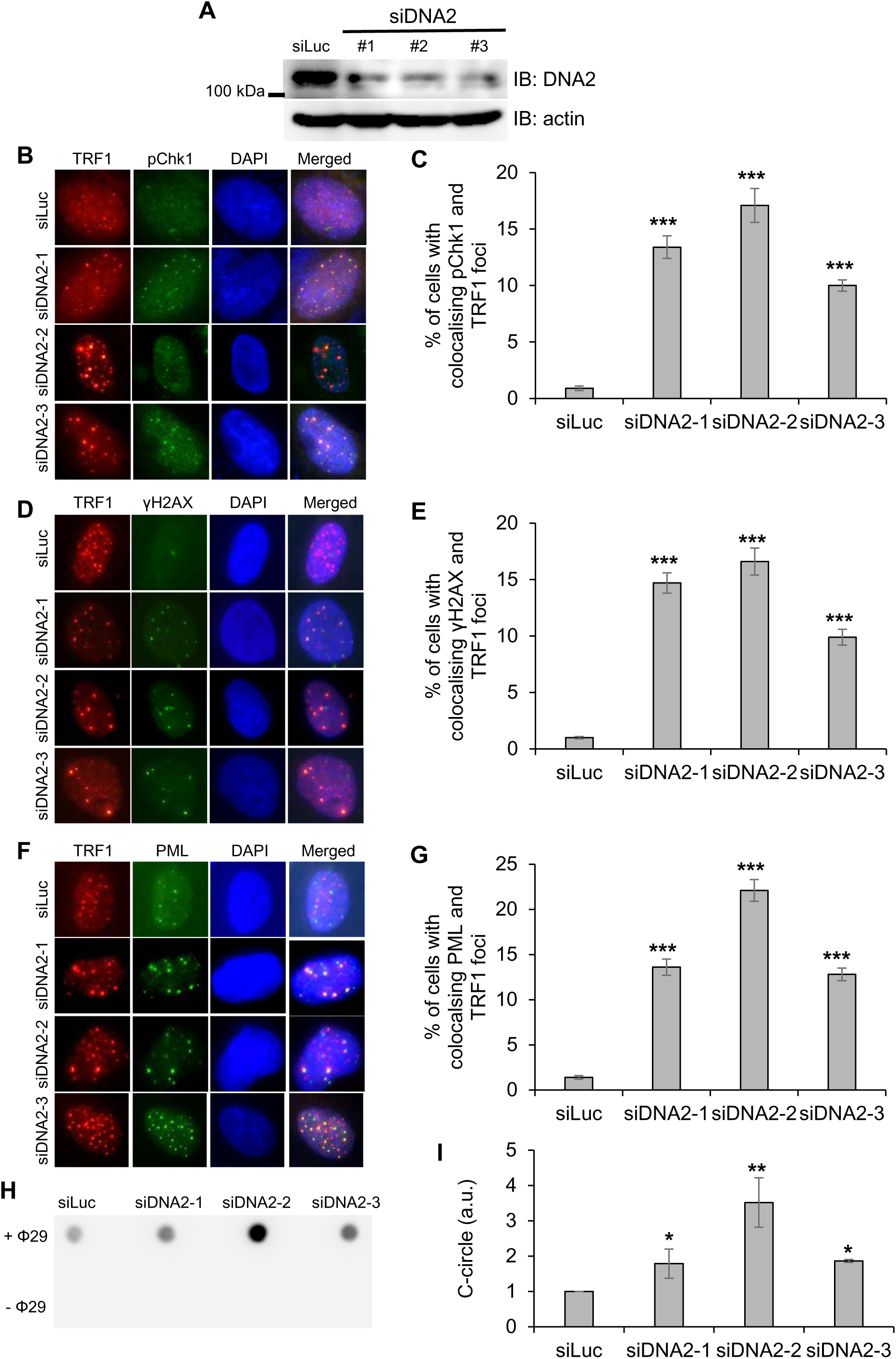
Depletion of DNA2 in ALT+ cells induces a pronounced replication stress response at telomeres and an increase of many ALT properties. U2OS cells were transfected with three different siRNA targeting DNA2 (siDNA2-1, siDNA2-2, and siDNA2-3). A siRNA targeting luciferase (siLuc) was used as the negative control. (**A**) Validation of the knocking down efficiency of the siDNA2 through Western Blot. (**B** to **G**) siRNA transfected U2OS cells were stained with antibodies recognizing TRF1 and phospho-Chk1 (Serine-345) (pChk1) (B and C), or TRF1 and γH2AX (D and E), or TRF1 and PML (F and G). All nuclei were stained with DAPI. More than two hundred cells were counted for each sample. (**H** and **I**) U2-OS cells were first transfected with different siRNA. DNA was then extracted and used for C-circle assay. “– Φ” indicates samples with no Phi(Φ)29 DNA polymerase added. Image J was used for quantification of the images. All error bars are standard deviations obtained from three different experiments. Standard two-sided t test: \**p*<0.05, \*\**p*<0.01, \*\*\**p*<0.001.

Previously, we and others showed that increased replication stress at ALT telomeres leads to the increase of many ALT properties [1–4]. Similarly, depletion of DNA2 in U2OS and Saos2 induced a pronounced increase of many ALT properties, including TIFs (Figs 1D, 1E, S4D, and S4E), APBs (Figs 1F, 1G, S4F, and S4G), and C-circles (Figs 1H and 1I). The DNA2 inhibitor, d16, induced a pronounced increase of TIFs in U2OS (Figs S2E and S2F). Notably, these effects were not observed in A549 and HeLa (data not shown). In our previous study, we demonstrated that BRCA1 and BLM are both recruited to telomeres and are required for replication stress response at telomeres in the FANCM deficient ALT cells [1]. Similarly, BLM is recruited to the ALT telomeres in C5 treated U2OS (Figs S3C and S3D). Depletion of BRCA1 or BLM attenuated replication stress response at telomeres in the DNA2 deficient U2OS (Figs S5A to S5C, S5E to 5G), suggesting that BRCA1 and BLM also function upstream of DNA2 in the ALT pathway. In addition, knocking down BRCA1 reduced localization of BLM to telomeres (Fig 5D), indicating that BRCA1 facilitates the recruitment of BLM to ALT telomeres to produce extensive single-stranded DNA (ssDNA) and induce robust replication stress response.

Taken together, our data indicates that DNA2 plays a crucial role in the ALT pathway by suppressing replication stress and DNA damage at telomeres. Since both d16 and C5 bind the DNA-binding motif of DNA2 that is shared by its nuclease and helicase [44, 45], both nuclease and helicase activities of DNA2 may be important for its function in the ALT pathway. When DNA2 is inhibited, BRCA1 then recruits BLM to damaged ALT telomeres, generates extensive ssDNA, and induces a robust replication stress response and DNA damage, which leads to the increase of many ALT properties.

### Depletion of DNA2 leads to an accumulation of TERRA R-loops and a decrease in DNA replication efficiency at ALT telomeres

As mentioned above, telomeres, due to their unique sequences and genomic features, are classified as DTRs. One of the main replication impediments at telomeres is TERRA R-loops, whose levels are much higher in ALT+ cells than in TEL+ cells [20]. Next, we examined whether the increased telomeric replication stress and DNA damage seen in the DNA2 deficient ALT+ cells were due to increased TERRA R-loops. We first performed immunofluorescence staining using the monoclonal antibody S9.6 to detect the DNA:RNA hybrid in siDNA2 transfected U2OS cells. In order to avoid detection of double-stranded RNA (dsRNA), all cells were pre-treated with RNase III to remove the dsRNAs before staining. Simultaneously, cells were stained with an antibody recognizing TRF2 and hybridized with a probe recognizing the TERRA RNA. As seen in Figs 2A to 2C, we observed a pronounced increase of S9.6 foci and TERRA foci that colocalize with TRF2 in the DNA2 deficient cells, suggesting that inhibition of DNA2 leads to TERRA R-loop accumulation at ALT telomeres. In addition, we also performed the DNA:RNA immunoprecipitation (DRIP) assay using the S9.6 antibody. Compared to the control siRNA transfected cells, the S9.6 antibody pulled down twice the amount of telomeric DNA:RNA hybrids in DNA2 depleted cells, which were eliminated when pre-treated with RNase H1, a nuclease that specifically degrades RNA within the DNA:RNA hybrids (Fig 2D). Collectively, our data demonstrated a pronounced increase of TERRA R-loops in the DNA2 deficient ALT+ cells compared to the control cells.

**Figure 2.**
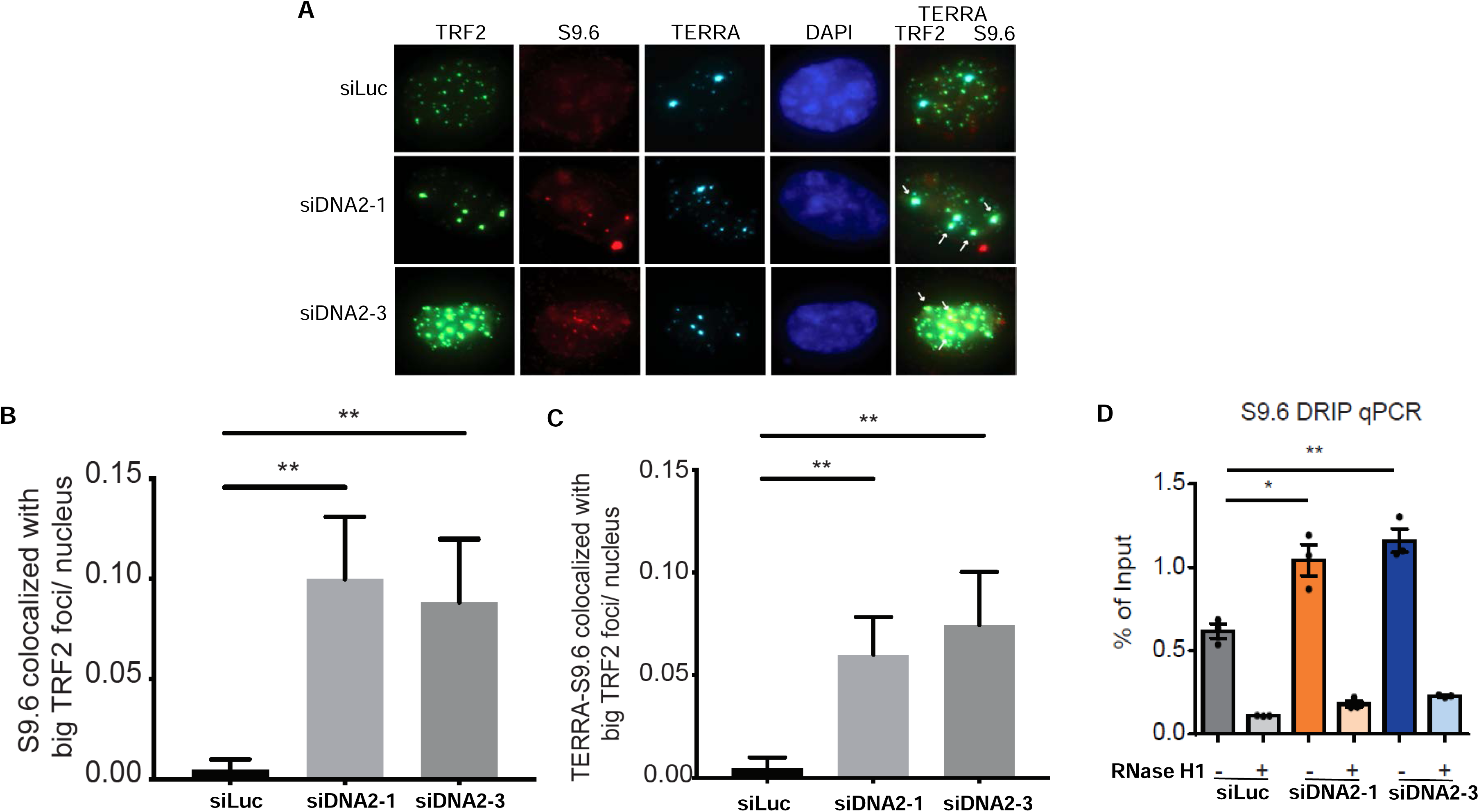
Depletion of DNA2 in ALT+ cells induces TERRA R-loop accumulation. U2OS cells were transfected with siRNA targeting either luciferase (siLuc) or DNA2 (siDNA2-1 and siDNA2-3). (**A** to **C**) Immuno-RNA FISH staining for TERRA, S9.6, and TRF2 were performed and quantified. The number of S9.6 foci that co-localize with the TRF2 foci was quantified in (B). The number of S9.6 foci that co-localize with the TRF2 foci and TERRA foci was quantified in (C). RNase III was added to remove double-stranded RNA signals prior to immune-RNA FISH for TERRA, S9.6, and TRF2. (**D**) siRNA transfected U2OS cells were used for S9.6 DRIP qPCR assay. All data were collected from two biological replicates. Standard two-tailed Student’s t-test: \**p*<0.05, \*\**p*<0.01, \*\*\**p*<0.001.

Next, we performed the single-molecule analysis of replicated DNA (SMARD) assay to determine whether the accumulated TERRA R-loops directly impede DNA replication at ALT telomeres [21]. Briefly, siRNA transfected U2OS cells were sequentially pulse-labeled with 5-iodo-2-deoxyuridine (IdU) for four hours and then 5-chloro-2-deoxyuridine (CldU) for four hours (Fig 3A). Genomic DNA was carefully extracted and digested with Pme I, which has a consensus sequence of GTTT/AAAC, in order to cut the genomic DNA but preserve the large DNA fragments. The digested genomic DNA was then separated by pulsed-field gel electrophoresis (PFGE), which is suitable for resolving the large DNA fragments. DNA fragments with sizes ranging from 160 kb to 200 kb and enriched subtelomeric and telomeric DNA were isolated and stretched on coverslips and then stained with antibodies recognizing IdU (red in Fig 3A) and CldU (green in Fig 3A). DNA molecules containing telomeres were identified by a fluorescent *in situ* hybridization (FISH) probe specific for telomeres (blue in Fig 3B). The telomere molecules with red signal, green signal, or both were identified as having undergone active and processive DNA replication during the eight-hour pulsed-label period (Fig 3C). In control siRNA transfected U2OS cells, we observed approximately 34% telomeres that underwent active replication, which is similar to what we reported previously [1]. In contrast, the actively replicated telomeres in siDNA2 transfected cells were reduced to approximately 21%, a 38% decrease, indicating that the accumulated TERRA R-loops observed in the DNA2 deficient ALT cells severely compromised the DNA replication efficiency at telomeres.

**Figure 3.**
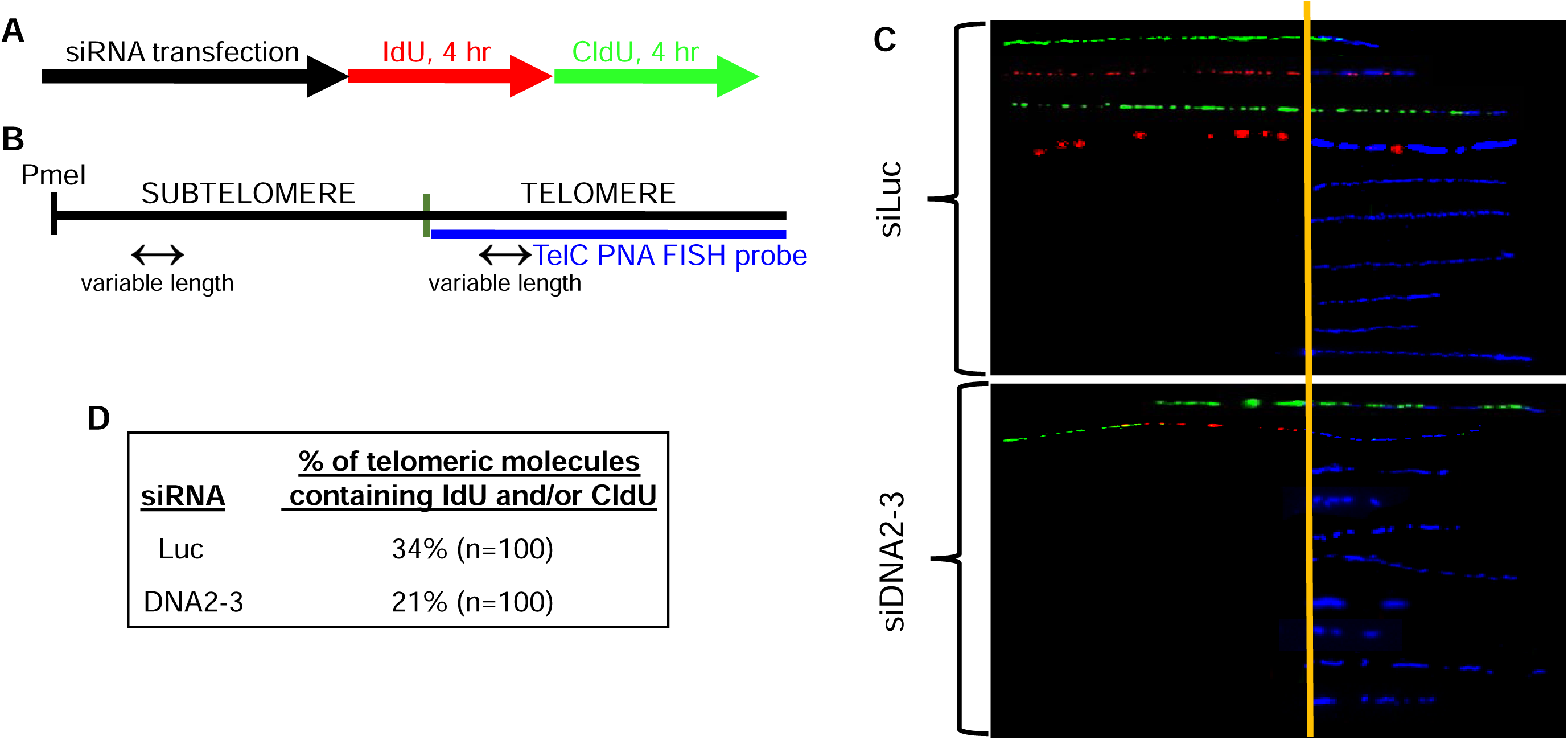
Depletion of DNA2 in ALT+ cells reduces DNA replication efficiency at telomeres. **(A** to **C**) U2OS cells were first transfected with siRNA targeting Luciferase control (siLuc) or DNA2 (siDNA2-3). 24 hours after the second siRNA transfection, cells were sequentially pulse-labeled with IdU for 4 hours and then CldU for 4 hours (A). SMARD was performed on telomeric DNA PmeI fragments isolated from these cells as detailed in the **Material and Methods**, with fragments ranging from 160-200 kb that contained subtelomeric DNA of variable lengths. The telomeric DNA molecules of variable lengths were identified by FISH using telomeric PNA probes (TelC; blue), while incorporated IdU and CldU were detected by indirect immunofluorescence with antibodies against IdU (red) and CldU (green). The diagram of the PmeI fragment including the subtelomeric and telomeric DNA and the representative images of subtelomeric/telomeric fragments are shown in (B). A few representative SMARD molecules are shown in (C). The vertical orange lines in (C) mark the boundary between the subtelomere and telomere. (**D**) The percentage of telomeric molecules that contained IdU and/or CldU (i.e., the ratio of labeled versus unlabeled fragments) is presented. This percentage indicates the relative efficiency of processive DNA replication in the regions of subtelomeres and telomeres during the IdU/CldU labeling period.

Taken together, our data showed that depletion of DNA2 induces a dramatic accumulation of TERRA R-loops at ALT telomeres, which then impedes active and processive DNA replication and induces robust replication stress response and DNA damage.

### DNA2 and FANCM manifest a strong additive genetic interaction in the ALT pathway

We and others established FANCM as one of the most important factors in the ALT pathway [1–4]. Next, we investigated the genetic interaction between DNA2 and FANCM in the ALT pathway. Excitingly, when we co-depleted DNA2 and FANCM in U2OS cells, we observed a strong additive effect in inducing replication stress response (Fig 4A), generating ssDNA (Fig 4B) and DNA damage (Fig 4C) at telomeres. Similar results were obtained in Saos2 cells (Figs S6A to S6C). Consistently, we observed an additive effect between DNA2 and FANCM in stimulating C-circle formation in both U2OS cells (Figs 4E and 4F) and Saos2 cells (Figs 6G and 6H). Intriguingly, co-depletion of DNA2 and FANCM did not recruit additional BLM to the ALT telomeres, suggesting that BLM protein may be limited in quantity in the ALT+ cells (Figs 4D and S6D).

**Figure 4.**
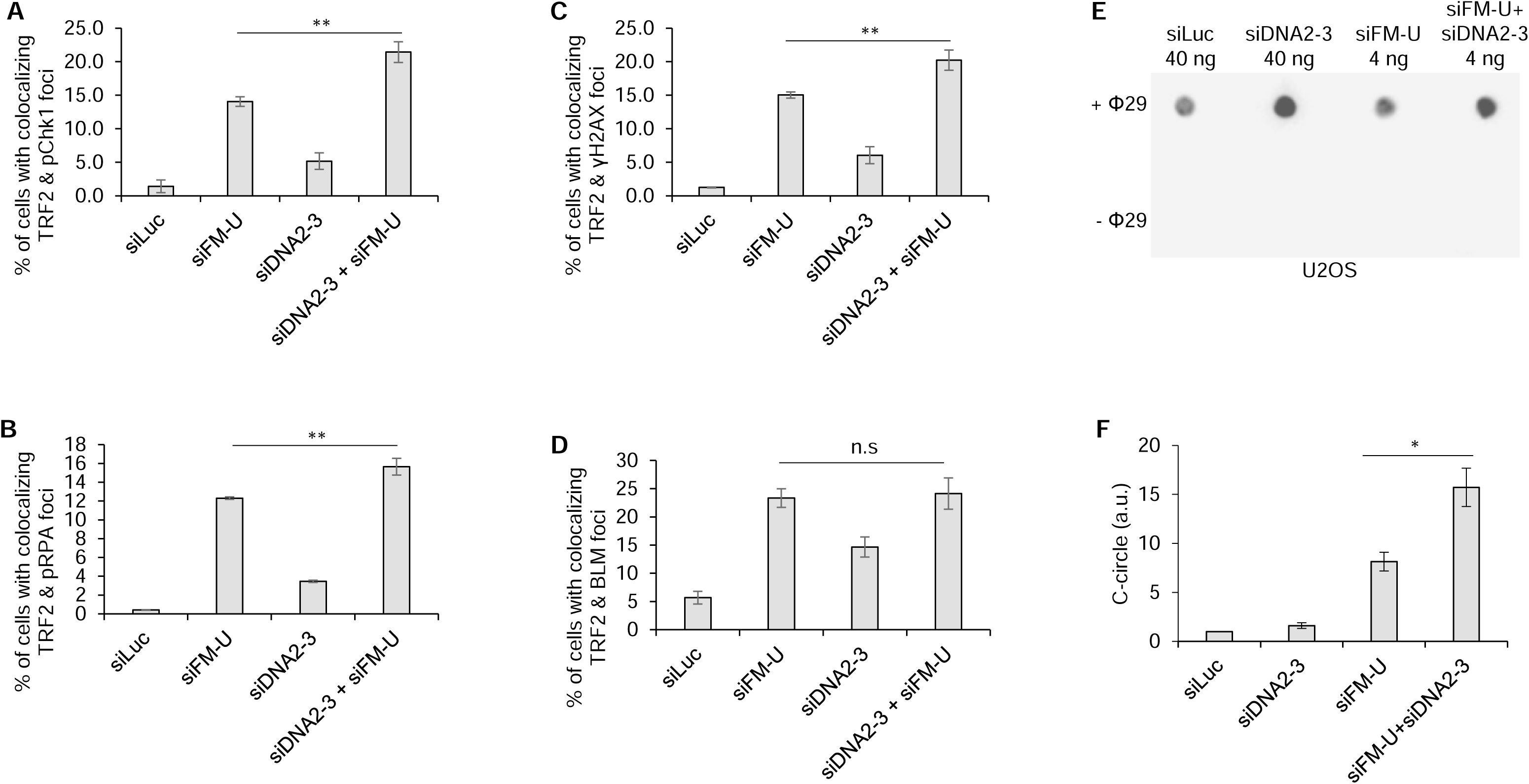
DNA2 and FANCM manifest a strong additive genetic interaction in suppressing replication stress at ALT telomeres. (**A** to **D**) U2-OS cells were first transfected with siRNA and then stained with antibodies recognizing TRF1 and pChk1 (A), or TRF1 and γH2AX (B), or TRF1 and pRPA (C, pRPA), or TRF1 and BLM (D). All nuclei were stained with DAPI. More than two hundred cells were counted for each sample. (**E** and **F**) C-circle assay. “–Φ” indicates samples with no Phi(Φ)29 DNA polymerase added. Image J was used for quantification of the images. All error bars are standard deviations obtained from three different experiments. Standard two-sided t test: \**p*<0.05, \*\**p*<0.01, \*\*\**p*<0.001. n.s., not significant.

Next, we performed the ATSA assay to determine the function of DNA2 and FANCM in break-induced telomere synthesis (BITS) [46]. Interestingly, we observed a robust increase of BITS in both DNA2 deficient cells and FANCM deficient cells, the large majority of which take place in APBs (Figs 5A and 5B). Consistent with the results shown above, in DNA2 and FANCM co-depleted cells, we observed a strong additive effect in BITS (Fig 5B), APB formation (Fig 5C), and telomere clustering (Fig 5D), another prominent ALT property that resulted from protein phase separation [47, 48]. Most importantly, we found that co-depletion of DNA2 and FANCM is synthetic lethal in the two ALT+ cells (Fig 5E), but not in the two TEL+ cells (Fig 5F).

**Figure 5.**
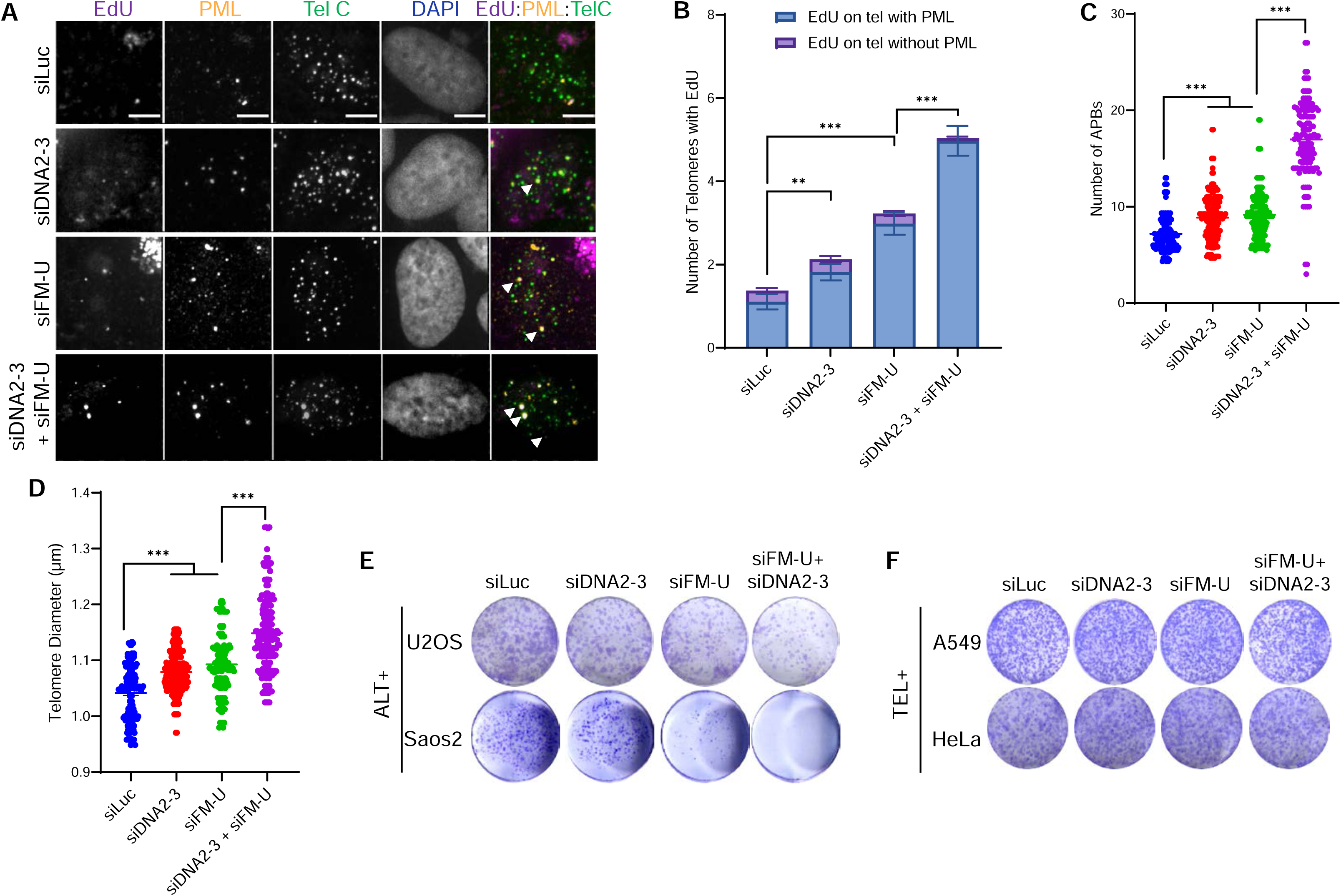
Co-depletion of DNA2 and FANCM induces robust break-induced telomere synthesis (BITS) and cause synthetic lethality in ALT+ cells. **(A and B).** Representative images (A) and quantification of EdU signal at telomeres (B) in U2OS. Arrow heads indicate some of the EdU signal at telomeres. Scale bars, 5 μm. **(C and D).** Quantification of the number of APBs (C) and the diameters of telomeres (D). Each dot represents one cell in three independent experiments, more than 70 cells were analyzed in each group. (**E** and **F**) Long-term viability assay. Two ALT+ cells (U2OS and Saos2) (E) and two TEL+ cells (A549 and HeLa) (F) were first transfected with different siRNA and then were used for crystal violet assay.

Taken together, our data strongly indicates that in ALT+ cells, DNA2 and FANCM function in two distinctive pathways to suppress replication stress response at telomeres by actively disrupting TERRA R-loops, thus prevent excessive DNA damage and promote their survival.

### DNA damage induced by the deficiency of DNA2 or FANCM stimulates telomere elongation

Recently, Xiao and colleagues developed the single-molecule telomere assay via optical mapping (SMTA-OM) to define genome-wide features of telomeres/chromosome ends at the single-molecule level [27, 49, 50]. Since DNA2 and FANCM play such a critical role in the ALT pathway, we utilized the SMTA-OM technology to thoroughly characterize the telomeres/chromosome ends in the DNA2 and FANCM deficient cells. Briefly, carefully extracted large genomic DNA molecules (>300 kb) are either labeled directly by a direct-labeling enzyme (DLE-1, Bionano Genomics), or nicked by a nickase Nt. BspQI (5’-GCTCTTCN-3’) first and then labeled. The motif sequences identified by DLE-1 and Nt. BspQI throughout the genome are labeled with a green fluorophore. Subsequently, telomeres are recognized and nicked with a telomere guide RNA (gTelo) and CRISPR/Cas9D10A and then also labeled with a green fluorophore [49, 51]. All DNA molecules are co-stained with YOYO-1 (blue). The two-color (green and blue) labeled DNA molecules are linearized in the NanoChannel Arrays and imaged. The patterns of spaced green labels from Nt. BspQI or DLE-1 are used to identify a specific chromosome arm by matching them to the predicted patterns of a reference genome. A more intense and uniform green signal at the end of a specific chromosome arm is identified as the end telomere (End Tel, Fig 6A). As reported previously, SMTA-OM can not only identify chromosome arm-specific end telomeres and measure their length, but it can also identify and quantify additional features at the chromosome ends, including two types of chromosome fusions, fusion/ITS+ (ITS+, Fig 6A) and fusion/ITS-(ITS-) [27, 50]. ITS+ manifests as a DNA molecule that on one side of the telomere signal, the DNA fragment is long enough to be assigned to a specific chromosome arm based on the pattern of its green labels. However, on the other side of the telomere signal, because the DNA fragment is too short, it cannot be confidently assigned to a particular chromosome arm based on the pattern of its green labels.

**Figure 6.**
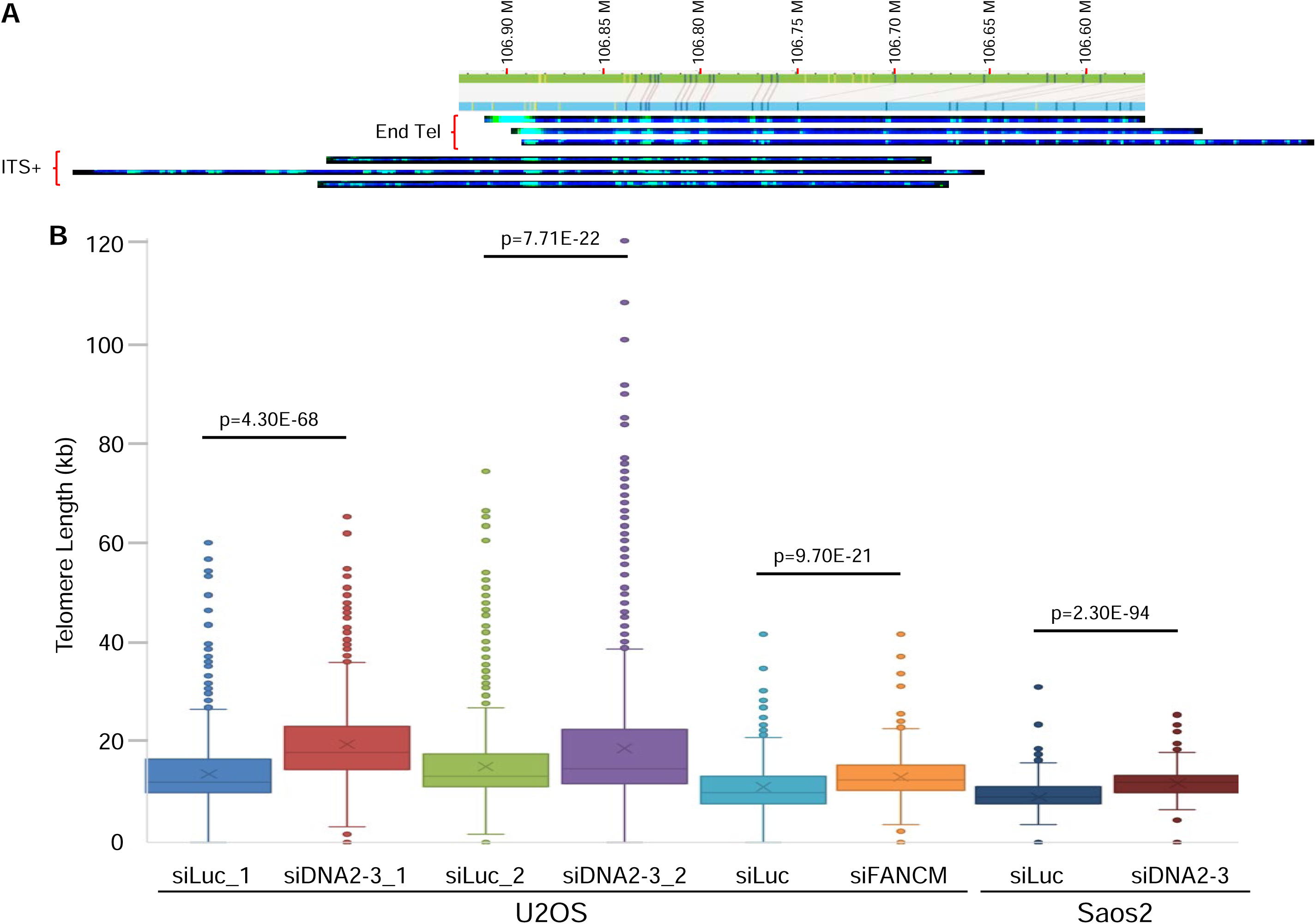
Depletion of DNA2 or FANCM increased the average end telomere (End Tel) length in ALT+ cells. U2OS or Saos2 cells were first transfected with siLuc, or siDNA2-3, or siFANCM (siFM-U) and then were used for SMTA-OM assay. (**A**) Representative images of DNA molecules assigned to chromosome 14q. (**B**) The average end telomere (End Tel) length from different siRNA transfected cells. P values are indicated on the top.

To demonstrate the reproducibility of the SMTA-OM, we performed the SMTA-OM assay using U2OS cells transfected with either siLuc or siDNA2-3 from two independent experiments. Fig 6A shows a few representative SMTA-OM molecules derived from chromosome 14q of the siDNA2-3 transfected U2OS cells. On the top panel, it shows three End Tel molecules with different telomere lengths. On the bottom panel, it shows three ITS+ molecules, in which the right parts of the molecules are assigned to chromosome 14q. All the DNA molecules detected and imaged for chromosome 14q from both siLuc and siDNA2 transfected U2OS cells of the second experiment are shown in Fig S7. On average, there are 35 chromosome arms that can be imaged and analyzed in both experiments with a total of more than fourteen hundred molecules for each sample (Table 1 and Supplemental Table 1). The number of molecules detected for each chromosome arm varies from 8 to 80. When considering all of the chromosome arms together, we observed a pronounced increase in End Tel mean length of siDNA2-3 transfected cells compared to siLuc transfected cells in both experiments, a 44% increase for the first experiment and a 25% increase for the second experiment (Fig 6B and Table 1).

**Table-1:** A summary of overall end telomere (End Tel) length change.

Next, we analyzed the End Tel mean length changes for an individual chromosome arm with 25 or more molecules (Fig 7A and Supplemental Table 1). Among the 22 arms analyzed, 21 arms (95%) showed a consistent increase in both experiments while only one arm, 15q, showed inconsistency (decreased by 9.7% in the first experiment while increased by 82% in the second experiment), indicating that overall, the SMTA-OM assay is highly reproducible. As for the ITS+ molecules, they can be detected for most chromosome arms but only account for a small percentage of the total SMTA-OM molecules (Supplemental Table 1). However, certain chromosome arms are more prone to produce ITS+ than others. For example, close to half of the molecules of chromosome arms 1q and 18q in siLuc transfected cells are ITS+. Intriguingly, depletion of DNA2 reduced the ITS+ of 1q (by 22% for the first experiment and by 8% for the second experiment), but increased the ITS+ of 18q (by 9% for the first experiment and by 20% for the second experiment), suggesting that DNA2 may have different function at 1q and 18q in regulating the formation of ITS+.

**Figure 7.**
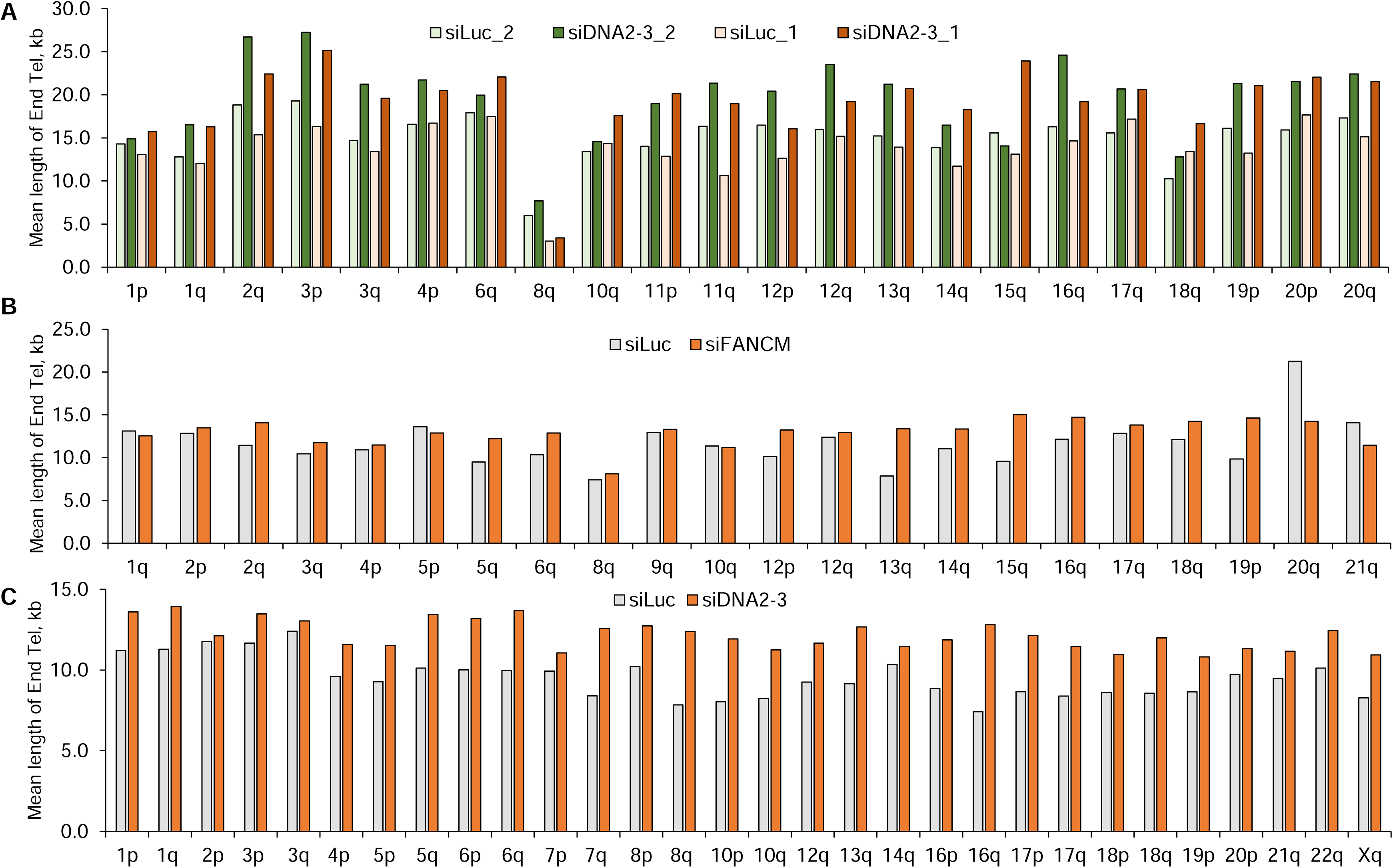
Genome-wide analysis of telomeres/chromosomes ends at individual chromosome arm level in DNA2 deficient cells and FANCM deficient cells. U2OS cells (**A** and **B**) or Saos2 (**C**) were first transfected with siLuc, or siDNA2-3, or siFANCM (siFM-U) and then were used for SMTA-OM assay. The mean length of the end telomeres (End Tel) was plotted for individual chromosome arms.

Next, we performed the SMTA-OM assay using Saos2 cells. Again, we observed a 25.8% increase of End Tel mean length in the siDNA2-3 transfected cells compared to the control cells (Fig 6B and Table 1). For the 30 chromosome arms with more than 26 molecules, all of them manifest increased End Tel mean length in the siDNA2-3 transfected cells compared to the control, ranging from a 3% increase for 2p to a striking 58% increase for 8q (Fig 7C). For the ITS+ molecules, 27 arms manifested an increase in the siDNA2-3 transfected cells compared to the control, while only three arms manifested a decrease (14q, 18p, and 20p) (Supplemental Table 1).

We also performed the SMTA-OM assay using U2OS cells transfected with either siLuc or siFANCM. Approximately 34 chromosome arms were imaged and analyzed with a total of more than one thousand molecules for each sample (Supplemental Table 1). When considering all chromosome arms together, we observed a 16.9% increase in End Tel mean length of siFANCM transfected cells compared to siLuc transfected cells (Fig 6B and Table 1). We also analyzed End Tel mean length changes for individual chromosome arm with more than 26 molecules (Fig 7B and Supplemental Table 1). Among the 22 arms analyzed, 17 arms (77%) manifest a consistent increase of End Tel mean length in siFANCM transfected cells compared to control cells while 5 arms (23%) showed a decrease. Most intriguingly, depletion of FANCM also reduced the ITS+ of 1q but increased the ITS+ of 18q, suggesting that FANCM and DNA2 may have similar functions at 1q and 18q in regulating the formation of ITS+.

Finally, we compared the SMTA-OM results in all four experiments to see if we could identify any common trend. Strikingly, five arms (6q, 12q, 13q, 16q, and 20p, highlighted in yellow in Supplemental Table 2) manifest an increase in both End Tel and ITS+ in siDNA2 or siFANCM transfected cells compared to siLuc transfected cells, suggesting that at least for these five arms, increased telomere length and ITS+ are likely regulated by the same mechanism.

Collectively, the SMTA-OM analysis suggests that telomeric damage seen in the DNA2 and FANCM deficient cells likely activate the BITS, which then promotes telomere elongation at most chromosome arms and induces the formation of ITS+ at certain specific arms.

## Discussion

Compared to the telomerase, the detailed mechanism of the ALT pathway warrants further investigation. Here, we identified DNA2 as a *bona fide* new player in the ALT pathway. We showed that depletion of DNA2 induces a robust replication stress response at ALT telomeres, leading to an increase in DNA damage and many ALT properties. Most importantly, we demonstrate that DNA2 suppresses replication stress at ALT telomeres by actively disrupting TERRA R-loops. Furthermore, we uncovered a strong additive genetic interaction between DNA2 and FANCM in the ALT pathway. Finally, using the recently developed SMTA-OM technology, we thoroughly characterized genome-wide features of telomeres/chromosome ends in the DNA2 and FANCM deficient ALT+ cells at the single-molecule level. We found that telomeric damage induced by DNA2 and FANCM deficiency activates the break-induced telomere synthesis and promotes telomere elongation at most chromosome arms. Most intriguingly, we found that DNA2 and FANCM may also play a role in regulating chromosome arm-specific changes at telomeres/chromosome ends. This is the first study that demonstrates the role of a specific protein in regulating chromosome arm-specific properties of telomeres. Collectively, our data indicate that at ALT telomeres, DNA2 and FANCM function in distinctive pathways to actively disrupt TERRA R-loops and thus prevent excessive replication stress and DNA damage, which help to balance the maintenance of telomere length and the survival in ALT+ cells.

### Chromosome arm-specific regulation of telomeres

A long-standing question in the field of telomere biology is whether telomeres at different chromosome ends are maintained and behave in the same way. Recent studies suggest that the answer is likely “NO”. For example, using an Oxford Nanopore Technology (ONT)-based long-read sequencing, called TeloTag, Karimian and colleagues showed that in a variety of non-transformed human cells, not only telomeres from different chromosome ends manifest different mean lengths, the differences are also present at birth and are maintained during the aging process [52]. In addition, they also found that for certain chromosome arms, for example 1p, 4q, and 21p, the mean lengths of telomere from the maternal haplotype are different from those of the paternal haplotype. In a separate study, also using the ONT-based long-read sequencing, called Telo-seq, Schmidt and colleagues observed similar chromosome arm-specific and haplotype-specific telomere properties in normal human cells [53]. Most recently, we also reported unique chromosome arm-specific changes in response to dCas9/sgTelo-induced replication stress at telomeres [54]. For example, we detected only 2.9% of chromosome 8q-containing fusion/ITS+ in control cells. In stark contrast, the chromosome 8q containing fusion/ITS+ in dCas9/sgTelo cells increased to 35.4%, a drastic 12-fold increase! In this study, we found that DNA2 and FANCM may regulate replication stress differently at different ALT telomeres. For example, depletion of FANCM in U2OS cells stimulates the elongation of telomeres at certain chromosome arms but not the others (Fig 7B and Supplemental Table 1). In comparison, depletion of DNA2 in both U2OS and Saos2 cells stimulates the elongation of telomeres in almost all chromosome arms (Figs 7A and 7C, and Supplemental Table 1). On the other hand, DNA2 and FANCM may also have certain overlapping functions at certain chromosome arms. For example, in both DNA2 and FANCM depleted U2OS cells, fusions/ITS+ of chromosome 1q are reduced, whereas fusions/ITS+ of chromosome18q are increased (Supplemental Table 1). This is the first study that establishes an important role of a particular protein in regulating properties of chromosome arm-specific telomeres.

The next logical question is what contributes to the chromosome arm-specific properties of telomeres. Accumulating evidence suggest that either the expression level of TERRA from different chromosome arms or the amount of TERRA R-loops formed at different chromosome arms, or both, may be implicated in regulating these chromosome arm-specific features [55–59]. For example, using RT-qPCR assay, Feretzaki and colleagues quantified the expression of TERRA from different chromosomes in different human cell lines and showed that: (1) the expression level of TERRA varies from chromosome arm to chromosome arm in the same cell line; (2) For the same chromosome arm, in most cases, ALT+ cells express higher levels of TERRA than TEL+ cells and non-transformed cells [59]. Most recently, combining biochemical enrichment of TERRA RNA and a ONT-base long-read sequencing, called TERRA ONTseq, Rodrigues and colleagues showed that: (1) TERRA RNA is expressed from most chromosome arms in human cells; (2) the amount of TERRA RNA expressed varies from chromosome arm to chromosome arm; (3) there are at least two different promoters that control the expression of TERRA RNA at different chromosome arms: the 29 bp repeat-related sequence and the 7q/12q promoter. At present, whether and how TERRA RNA or TERRA R-loops, or both, regulate the chromosome arm-specific properties remains a mystery. However, since both DNA2 and FANCM can regulate the formation of TERRA R-loops at ALT telomeres (Fig 2)[2, 4], we speculate that they may exert their chromosome arm-specific function at telomeres through regulating the level of TERRA R-loops. Uncovering the detailed mechanism behind these chromosome arm-specific regulations of telomeres/chromosome ends will be the next frontier of telomere research and may have important implications for human health.

### TERRA R-loops: a double-edged sword for the survival of ALT+ cells

As mentioned above, telomeres are DTR genomic loci, which pose challenges to the replisome and are more prone to DNA damage. TERRA R-loops are likely the primary DNA replication impediments at telomeres, especially in ALT+ cells [60, 61]. Cesare and colleagues showed that overall spontaneous telomeric DNA damage is much higher in ALT+ cells than in TEL+ cells [33]. Increased expression of TERRA and/or accumulation of TERRA R-loops are the likely causes of heightened spontaneous telomeric DNA damage in ALT+ cells [20, 62, 63]. In the absence of telomerase, mildly increased spontaneous telomeric DNA damage may be beneficial for ALT+ cells because they can activate the break-induced telomere synthesis (BITS), which then facilitates the elongation of the short telomeres and thus avoid excessive telomere shortening induced cell death. Indeed, we observed an increase in the average telomere length in both DNA2 and FANCM deficient ALT+ cells (Table 1 and Fig 6). However, previous studies and the data presented here indicate that too many TERRA R-loops can be detrimental to the survival of ALT+ cells because they will overwhelm the ALT+ cells with excessive replication stress and DNA damage at their telomeres, eventually leading to reduced viability (Fig 5E) [2, 4]. Therefore, ALT+ cells need to fine-tune the amount of TERRA transcribed at their telomeres and prevent the excessive amount of TERRA R-loops by recruiting DNA2, FANCM, RNase H1, and other proteins to proactively disrupt the TERRA R-loops [2, 4, 20, 61].

### Targeting the replication stress response for treating ALT+ cancer

Unlike many TEL+ cancers, chemotherapy remains the main treatment option for most ALT+ cancers [14]. Previously, we and others have shown that accumulation of excessive amounts of TERRA R-loops could be the Achille’s heel of the ALT+ cells [1, 3, 4]. Here we showed that co-depletion of DNA2 and FANCM induce synthetic lethality in the two ALT+ cells but not in the two TEL+ (Figs 5E and 5F). Therefore, both FANCM and DNA2 are potential drug targets for treating ALT+ cancer.

## Acknowledgments

We want to thank Liam Buchanan for proofreading the manuscript.

## Author contributions

Conceptualization: D.Z. and M.X. Methodology and investigation: A.R., H.Z.A., Y.C., R.Z., S.T.K., D.I.Y., J.N., R.F., S.T., B.Y., M.S., L.Z., Writing, review & editing: D.Z., B.S., H.Z., H.P.C.C., C.L.S., M.X.

## Materials and Methods

### Cell lines, tissue culture, and chemicals

A549, HeLa, U2OS and Saos2 were purchased from ATCC. All cells are grown in DMEM (Corning) supplemented with 10% fetal bovine serum (Bio-techne), penicillin (Corning), and streptomycin (Corning), at 37 °C in a humidified incubator with 5% CO_2._ C5 was generously provided by Dr. Binghui Shen (City of Hope). D16 was purchased from Molport Inc.

### siRNA transfection

Cells are transfected twice on consecutive days using 50 nM siRNA with RNAiMAX (Invitrogen) following the protocol recommended by the manufacturer. The sequence and order information for all the siRNA are listed in the Supplemental Table-3.

### Immunoblotting

Cells are collected and lysed directly in a sample buffer, followed by sonication for 30 seconds. Equal volumes of lysate were loaded for immunoblotting. The DNA2 antibody was generously provided by Dr. Binghui Shen (City of Hope). The Actin antibody was purchased from Santa Cruz (sc-1616).

### Immunofluorescent staining

Cells were seeded onto coverslips. Cells were either then fixed with 3% paraformaldehyde containing 2% sucrose for 10 min, followed by treatment with Triton X-100 solution on ice for 5 min, or treated with Triton X-100 solution on ice for 5 min then fixed with 3% paraformaldehyde containing 2% sucrose for 10 min (pre-extraction). Slides were then blocked with 1% Gelatin in PBS. Cells are stained with primary antibodies and the respective Alexa-488 (Invitrogen) and Alexa-546 (Invitrogen) conjugated secondary antibodies. All the antibodies used for immunofluorescent staining are listed in the Supplemental Table-4. SlowFade Gold DAPI (S36938) was used to stain the DNA. Images were collected by using an Olympus upright Fluorescent Microscope images were processed using Adobe Photoshop.

### C-circle assay

Cells were transfected twice with different siRNA. 48 hours later, genomic DNA from 3-5 x 10^5^ cells was extracted using QIAamp DNA Blood Mini Kit (Qiagen, 51106). 4 μg genomic DNA were digested with Alu I (NEB, R0137L) and Mbo I (NEB, R0147L) at 37 °C for at least 2 hours and then purified using Qiagen PCR Purification Kit (Qiagen, 28106). 40 ng of Alu I and Mbo I digested DNA were used for the Phi29 DNA polymerase reaction (30 °C for 8 hours and then 65 °C for 20 min). All the PCR reaction mixtures were loaded onto the Amersham Hybond-N+ membrane using the Bio-Rad Bio-Dot Microfiltration apparatus. The membrane was cross-linked using the UV Stratalinker at energy 1200. The membrane was then processed, probed, and developed using the TeloTAGGG Telomere Length Assay kit (Roche, 12-209-136-001). Images were taken with an Amersham Imager 600 and quantified using the NIH ImageJ.

### Immuno-RNA-FISH

U2OS cells grown on cover slips were washed with cold PBS and treated with CSKT (10 mM PIPES, pH 6.8, 100 mM NaCl, 3 mM MgCl_2_, 0.3 M sucrose, 0.5% Triton X-100, adjust to pH 6.8) for 10 minutes on ice. Cells were fixed in 4 % paraformaldehyde at RT and stored at 70% ethanol at -20°C. After washing with cold PBS, cells were incubated with blocking solution (1% BSA/PBS with 1 mM EDTA and 0.8 U/μl of RNase inhibitor) at 4°C for 1 hr. Cells were then incubated with primary antibodies in blocking solution at 4°C overnight, and washed with 0.2% Tween20/PBS three times at 4°C. Antibodies were used against TRF2 (Novus, NB110-57130) and S9.6 (1:100 dilution, Millipore, # MABE1095). Followed by incubation of secondary antibodies (in blocking solution) at 4°C for 2 hours, cells were washed with PBS three times and fixed in 2 % paraformaldehyde for 10 minutes. RNA FISH was then performed after immunostaining. TERRA oligo probes ((TAACCC)_7_-Alexa-647-3’) for RNA-FISH were mixed at the final concentration of 0.5 pmol/μl in hybridization solution (50% formamide, 2×SSC, 2 mg/ml BSA, 10% Dextran Sulfate-500K). Hybridization was carried out at 42°C overnight for RNA FISH. Cells were washed with 2×SSC/50% formamide for 5 min three times at 44°C, and then washed with 2×SSC for 5 min twice at 44°C. Images were captured using Olympus IX83 inverted microscopy with various Z-sections and then were compiled to 3D images to calculate APB and TERRA-associated APB foci. ABP foci were determined by the extensive TRF2 staining (> 3-fold increase compared to the average signal) with PML staining.

### DRIP-qPCR

DRIP protocol was conducted as described previously (PMID36184605). Briefly, cells were resuspended in nuclear extraction buffer (0.5% NP-40, 80 mM KCl, 0.5 mM HEPES pH 8.0) on ice for 30 min, and nuclei were spun down and resuspended in lysis buffer (1% SDS, 25 mM Tris-HCl, pH 8.0, 5 mM EDTA). The cell lysates were treated with proteinase K at 55°C for 3 hours. Genomic DNA was extracted by phenol-chloroform isoamyl alcohol (25:24:1) with the phase lock gel tubes and precipitated with ethanol. Genomic DNA was fragmented into 200∼500 bp using Covaris S2 in microtubes. For RNaseH controls, 8 μg of genomic DNA was treated with RNase H (5 U/μL, NEB, # M0523) at 37°C overnight and purified by phenol-chloroform extraction. For immunoprecipitation, 5 μg genomic DNA was used in 500 μL 1X binding buffer (10 mM sodium phosphate, pH 7, 140 mM NaCl, and 0.05% (vol/vol) Triton X-100) per capture and incubated with 2 μg of S9.6 antibody (Millipore, # MABE1095) at 4°C for overnight. 20 μL Protein G Dynabeads (Thermo Fisher Scientific, #10004D) was added and incubated at 4°C for 2 hours. After washes with 1X binding buffer for 15 min twice at RT, captured DNA was eluted in 300 μL elution buffer (50 mM Tris, pH 8.0, 10 mM EDTA, pH 8.0, 0.5% (vol/vol) SDS) containing 7 μL of 20 mg/mL proteinase K at 55°C for 45 min. DNA was purified by phenol-chloroform extraction and telomeric repeat DNA was measured by qPCR (forward primer: 5’-GGTTTTTGAGGGTGAGGGTGAGGGTGAGGGTGAGGGT-3’, reverse primer: 5’TCCCGACTATCCCTATCCCTATCCCTATCCCTATCCCTA-3’).

### Single Molecule Analysis of Replicated DNA (SMARD)

The SMARD assay was performed essentially as described previously[1]. Briefly, siRNA transfected U2OS cells were sequentially labeled with 30 μM IdU (4 hours) and 30 μM CldU (4 hours). DNA isolation and processing for SMARD were as described previously[1]. Following Pme I digestion, telomeric DNA fragments ranging from 160 to 200 kb were resolved by PFGE, identified by Southern blot, and then isolated. The Pme I-digested DNA was stretched on microscope slides coated with 3-aminopropyltriethoxysilane (Sigma). After stretching, the DNA was denatured in alkali-denaturing buffer (0.1 N NaOH in 70% ethanol and 0.1% β-mercaptoethanol) for 12 min and fixed by addition of 0.5% glutaraldehyde for 5 min. Telomeric DNA was identified by hybridization with a Biotin-OO-(CCCTAA)4 PNA probe (TelC, Biosynthesis) followed by incubation with Alexa Fluor 350-conjugated NeutrAvidin (Molecular Probes), followed by two rounds of incubation first with a biotinylated anti-avidin antibody (Vector) and then with the Alexa Fluor 350-conjugated NeutrAvidin. Incorporated halogenated nucleotides were detected with a mouse anti-IdU monoclonal antibody (BD) and a rat anti-CldU monoclonal antibody (Accurate) followed by Alexa Fluor 568-conjugated goat anti-mouse (Invitrogen) and Alexa Fluor 488-conjugated goat anti-rat secondary antibodies (Invitrogen). An antibody recognizing the single-stranded DNA was purchased from Millipore (anti-DNA antibody, clone 16–19, MAB3034).

### ALT telomere DNA synthesis in APBs (ATSA) assay

48 hours after the second siRNA transfection, cells were pulsed with EdU (10 μM) for 2 hours before harvest. Cells on glass coverslips were washed twice in PBS and fixed with 4% paraformaldehyde (PFA) for 10 min. Cells were permeabilized with 0.3% (v/v) Triton X-100 for 5 min. The Click-IT Plus EdU Cell Proliferation Kit with Alexa Flour 647 (C10635 Invitrogen) was applied to cells for 30 minutes to detect EdU.

Cells were then incubated with primary antibody at 4°C in a humidified chamber overnight and then with secondary antibody for 1 h at room temperature before washing and mounting. Primary antibodies were anti-PML (sc966, Santa Cruz, 1:100 dilution).

For IF-FISH, coverslips were first stained with primary and secondary antibodies, then fixed again in 4% formaldehyde for 10 min at room temperature. Coverslips were then dehydrated in an ethanol series (70%, 80%, 90%, 2 min each) and incubated with a TelC-Alexa488 probe (F1004, Panagene) at 75°C for 5 min and then overnight in a humidified chamber at room temperature. Coverslips were then washed and mounted for imaging.

Imaging acquisition was performed as previously described [64]. For fixed cell imaging, cells were seeded on 12 mm circular cover glass coverslips coated with poly-D-lysine (P1024, Sigma-Aldrich). Images were acquired with a microscope (ECLIPSE Ti2) with a 100 × 1.4 NA objective, a 16 XY Piezo-Z stage (Nikon Instruments Inc.), a spinning disk (Yokogawa), an electron multiplier charge-coupled device camera (IXON-L-897), and a laser merge module that was equipped with 488 nm, 561 nm, 594 nm, and 630 nm lasers controlled by NIS-Elements Advanced Research. Images were taken with 0.5 μm spacing between Z slices, for a total of 8 μm.

Images were processed and analyzed using NIS-Elements AR (Nikon). Maximum projections were created from z stacks, and thresholds were applied to the resulting 2D images to segment and identify telomere/PML foci as binaries. For quantification of the co-localization of two fluorescent labels, images were analyzed using binary operations in NIS-Elements AR. Co-localized foci were counted if the objects from different layers contained overlapping pixels.

### Crystal violet assay

4000 cells/per well are seeded in 12 well plates. After 7-14 days of growth, cells are fixed using a methanol/acetic acid solution and stained with 1% crystal violet solution.

### Single-molecule telomere assay via optical mapping (SMTA-OM)

Cells were transfected with different siRNAs, and stored in 10% DMSO in liquid nitrogen until DNA extraction and Purification.

#### DNA extraction and purification

Cells were first embedded in gel plugs with approximately 1 million cells per gel plug (BioRad no. 170-3592). The high molecular weight genomic DNA was extracted and purified using the Bionano Genomics SP kit.

#### DNA Labeling

Two motif-mapping approaches were used for the first part of the DNA labeling procedure. (1) The Nt.BspQI protocol, which was used for U2OS cells: The DNA sample was labeled as previously described [49]. (2) The DLE-1 protocol, which was used only for the Saos2 cells: Following the manufacturer’s instructions, a DLS labeling kit (Bionano Genomics) labeled 750 ng of genomic DNA. A labeling mix comprised Direct Labeling Enzyme 1 (DLE-1), 1X DLS reaction buffer, and DL green fluorophore-labeled nucleotide mix. The labeling mix was added to the genomic DNA, gently mixed via a wide bore pipette, and then incubated at 37°C for 2 hours. Following the incubation, membrane dialysis was used to remove the unwanted fluorescent dyes, proteins, and salts. Membrane dialysis was done at room temperature for approximately 2 hours in the dark to protect it from the light. Dialyzed DNA was then recovered using a 100 nm hydrophilic membrane (EMD Millipore, VCWP04700) and quantified with a Qubit device.

The second part of the DNA labeling procedure focuses on labeling the telomeres. Guide RNA (gRNA) consisting of 0.5 μM tracrRNA (IDT) and 50 μM crRNA was gently mixed via pipetting up and down and annealed on ice for 30 minutes. 25 pmol gRNA was incubated with 200 ng Cas9D10A nicking enzyme and 1X NE Buffer 3.1 at 37°C for 15 minutes. Next, 300 ng of the previously DLE-1 labeled genomic DNA was added to the Cas9D10A mixture for the one-hour nicking reaction at 37°C. After nicking, 200 nM red fluorophore-labeled nucleotides (ATTO647-dUTP, dATP, dGTP, dCTP) were incorporated into telomeres by 5U Taq DNA polymerase at 72°C for one hour in 1X Thermopol buffer (New England Biolabs). The nick-labeled sample was treated with proteinase K (QIAGEN) at 50°C for 30 min. Finally, to prepare the DNA for nanochannels, a staining mix of flow buffer, DTT, and YOYO-1 (a blue-colored dye that labels the backbone of DNA, Bionano Genomics DLS kit) was prepared according to the manufacturer’s instruction before adding it to the sample. The sample combined with the staining mix was incubated at room temperature overnight.

#### Imaging with Saphyr

A Bionano Saphyr G1.2 chip was loaded with the labeled DNA sample and imaged using a “dual-labeled sample” scheme incorporated in the Saphyr software. Images were taken in the following order: first, the red fluorescent labeling with the 637 nm laser, then the green fluorescent labeling with the 532 nm laser, and finally, the blue fluorescent DNA backbone YOYO-1 staining with the 473 nm laser.

#### Genome Assembly

*De novo* genome assembly was done using the software developed by Bionano Genomics. Consensus maps were generated by de novo assembling individual DNA molecules followed by alignment to the hg38 human reference genome.

#### Telomere analysis

CMAP, XMAP, and BNX files were generated after *de novo* genome assembly. The CMAP files contain both the red and the green label information. The XMAP files contain only the information on the green labels. Molecules matching the expected green labeling patterns were extracted from the BNX and CMAP files. Raw molecule images were extracted using in-house software based on their locations in the nanochannels. The red telomere labels were easily distinguishable from the green labels in the subtelomeric region. Telomeres were then analyzed and measured using the ImageJ software. The ferret diameter tool was used to capture the length of telomeres in pixels and intensities before converting all the measurements to kilobase (kb) following the previously established procedures[49, 50].

## References

1. Pan, X., et al., FANCM, BRCA1, and BLM cooperatively resolve the replication stress at the ALT telomeres. Proc Natl Acad Sci U S A, 2017. 114(29): p. E5940–E5949.

2. Pan, X., et al., FANCM suppresses DNA replication stress at ALT telomeres by disrupting TERRA R-loops. Sci Rep, 2019. 9(1): p. 19110.

3. Lu, R., et al., The FANCM-BLM-TOP3A-RMI complex suppresses alternative lengthening of telomeres (ALT). Nat Commun, 2019. 10(1): p. 2252.

4. Silva, B., et al., FANCM limits ALT activity by restricting telomeric replication stress induced by deregulated BLM and R-loops. Nat Commun, 2019. 10(1): p. 2253.

5. Ragupathi, A., et al., Targeting the BRCA1/2 deficient cancer with PARP inhibitors: Clinical outcomes and mechanistic insights. Front Cell Dev Biol, 2023. 11: p. 1133472.

6. Gadaleta, M.C. and E. Noguchi, Regulation of DNA Replication through Natural Impediments in the Eukaryotic Genome. Genes (Basel), 2017. 8(3).

7. Lezaja, A. and M. Altmeyer, Dealing with DNA lesions: When one cell cycle is not enough. Curr Opin Cell Biol, 2021. 70: p. 27–36.

8. Berti, M. and A. Vindigni, Replication stress: getting back on track. Nat Struct Mol Biol, 2016. 23(2): p. 103–9.

9. Zeman, M.K. and K.A. Cimprich, Causes and consequences of replication stress. Nat Cell Biol, 2014. 16(1): p. 2–9.

10. Cortez, D., Preventing replication fork collapse to maintain genome integrity. DNA Repair (Amst), 2015. 32: p. 149–57.

11. Marechal, A. and L. Zou, DNA damage sensing by the ATM and ATR kinases. Cold Spring Harb Perspect Biol, 2013. 5(9).

12. Lu, R. and H.A. Pickett, Telomeric replication stress: the beginning and the end for alternative lengthening of telomeres cancers. Open Biol, 2022. 12(3): p. 220011.

13. Cesare, A.J. and R.R. Reddel, Alternative lengthening of telomeres: models, mechanisms and implications. Nat Rev Genet, 2010. 11(5): p. 319–30.

14. MacKenzie, D., Jr., et al., ALT Positivity in Human Cancers: Prevalence and Clinical Insights. Cancers (Basel), 2021. 13(10).

15. Shay, J.W. and W.E. Wright, Telomeres and telomerase: three decades of progress. Nat Rev Genet, 2019. 20(5): p. 299–309.

16. Biffi, G., et al., Quantitative visualization of DNA G-quadruplex structures in human cells. Nat Chem, 2013. 5(3): p. 182–6.

17. Lam, E.Y., et al., G-quadruplex structures are stable and detectable in human genomic DNA. Nat Commun, 2013. 4: p. 1796.

18. Azzalin, C.M., et al., Telomeric repeat containing RNA and RNA surveillance factors at mammalian chromosome ends. Science, 2007. 318(5851): p. 798–801.

19. Schoeftner, S. and M.A. Blasco, Developmentally regulated transcription of mammalian telomeres by DNA-dependent RNA polymerase II. Nat Cell Biol, 2008. 10(2): p. 228–36.

20. Arora, R., et al., RNaseH1 regulates TERRA-telomeric DNA hybrids and telomere maintenance in ALT tumour cells. Nat Commun, 2014. 5: p. 5220.

21. Drosopoulos, W.C., et al., Human telomeres replicate using chromosome-specific, rather than universal, replication programs. J Cell Biol, 2012. 197(2): p. 253–66.

22. Sfeir, A., et al., Mammalian telomeres resemble fragile sites and require TRF1 for efficient replication. Cell, 2009. 138(1): p. 90–103.

23. Glover, T.W., T.E. Wilson, and M.F. Arlt, Fragile sites in cancer: more than meets the eye. Nat Rev Cancer, 2017. 17(8): p. 489–501.

24. Li, S. and X. Wu, Common fragile sites: protection and repair. Cell Biosci, 2020. 10: p. 29.

25. Bryan, T.M., et al., Telomere elongation in immortal human cells without detectable telomerase activity. EMBO J, 1995. 14(17): p. 4240–8.

26. Bryan, T.M., et al., Evidence for an alternative mechanism for maintaining telomere length in human tumors and tumor-derived cell lines. Nat Med, 1997. 3(11): p. 1271–4.

27. Raseley, K., et al., Single-Molecule Telomere Assay via Optical Mapping (SMTA-OM) Can Potentially Define the ALT Positivity of Cancer. Genes, 2023. 14(6): p. 1278.

28. Blasco, M.A., The epigenetic regulation of mammalian telomeres. Nat Rev Genet, 2007. 8(4): p. 299–309.

29. Doksani, Y., et al., Super-resolution fluorescence imaging of telomeres reveals TRF2-dependent T-loop formation. Cell, 2013. 155(2): p. 345–56.

30. Griffith, J.D., et al., Mammalian telomeres end in a large duplex loop. Cell, 1999. 97(4): p. 503–14.

31. de Lange, T., Shelterin: the protein complex that shapes and safeguards human telomeres. Genes Dev, 2005. 19(18): p. 2100–10.

32. Pfeiffer, V. and J. Lingner, Replication of telomeres and the regulation of telomerase. Cold Spring Harb Perspect Biol, 2013. 5(5): p. a010405.

33. Cesare, A.J., et al., Spontaneous occurrence of telomeric DNA damage response in the absence of chromosome fusions. Nat Struct Mol Biol, 2009. 16(12): p. 1244–51.

34. Suram, A., et al., Oncogene-induced telomere dysfunction enforces cellular senescence in human cancer precursor lesions. EMBO J, 2012. 31(13): p. 2839–51.

35. Sobinoff, A.P. and H.A. Pickett, Alternative Lengthening of Telomeres: DNA Repair Pathways Converge. Trends Genet, 2017.

36. Apte, M.S. and J.P. Cooper, Life and cancer without telomerase: ALT and other strategies for making sure ends (don’t) meet. Crit Rev Biochem Mol Biol, 2017. 52(1): p. 57–73.

37. Lee, M., et al., Telomere sequence content can be used to determine ALT activity in tumours. Nucleic Acids Res, 2018. 46(10): p. 4903–4918.

38. Gao, J. and H.A. Pickett, Targeting telomeres: advances in telomere maintenance mechanism-specific cancer therapies. Nat Rev Cancer, 2022. 22(9): p. 515–532.

39. Zheng, L., et al., Multiple roles of DNA2 nuclease/helicase in DNA metabolism, genome stability and human diseases. Nucleic Acids Res, 2020. 48(1): p. 16–35.

40. Hudson, J.J.R. and U. Rass, DNA2 in Chromosome Stability and Cell Survival-Is It All about Replication Forks? Int J Mol Sci, 2021. 22(8).

41. Choe, W., et al., Dynamic localization of an Okazaki fragment processing protein suggests a novel role in telomere replication. Mol Cell Biol, 2002. 22(12): p. 4202–17.

42. Lin, W., et al., Mammalian DNA2 helicase/nuclease cleaves G-quadruplex DNA and is required for telomere integrity. EMBO J, 2013. 32(10): p. 1425–39.

43. O’Sullivan, R.J., et al., Rapid induction of alternative lengthening of telomeres by depletion of the histone chaperone ASF1. Nat Struct Mol Biol, 2014. 21(2): p. 167–74.

44. Liu, W., et al., A Selective Small Molecule DNA2 Inhibitor for Sensitization of Human Cancer Cells to Chemotherapy. EBioMedicine, 2016. 6: p. 73–86.

45. Folly-Kossi, H., et al., DNA2 Nuclease Inhibition Confers Synthetic Lethality in Cancers with Mutant p53 and Synergizes with PARP Inhibitors. Cancer Res Commun, 2023. 3(10): p. 2096–2112.

46. Zhang, J.M., et al., Alternative Lengthening of Telomeres through Two Distinct Break-Induced Replication Pathways. Cell Rep, 2019. 26(4): p. 955–968 e3.

47. Zhang, H., et al., Nuclear body phase separation drives telomere clustering in ALT cancer cells. Mol Biol Cell, 2020. 31(18): p. 2048–2056.

48. Zhao, R., et al., SUMO promotes DNA repair protein collaboration to support alternative telomere lengthening in the absence of PML. Genes Dev, 2024. 38(13-14): p. 614–630.

49. McCaffrey, J., et al., High-throughput single-molecule telomere characterization. Genome Res, 2017. 27(11): p. 1904–1915.

50. Abid, H.Z., et al., Single-molecule analysis of subtelomeres and telomeres in Alternative Lengthening of Telomeres (ALT) cells. BMC Genomics, 2020. 21(1): p. 485.

51. McCaffrey, J., et al., CRISPR-CAS9 D10A nickase target-specific fluorescent labeling of double strand DNA for whole genome mapping and structural variation analysis. Nucleic Acids Res, 2016. 44(2): p. e11.

52. Karimian, K., et al., Human telomere length is chromosome end-specific and conserved across individuals. Science, 2024: p. eado0431.

53. Schmidt, T.T., et al., High resolution long-read telomere sequencing reveals dynamic mechanisms in aging and cancer. Nat Commun, 2024. 15(1): p. 5149.

54. Singh, M., et al., Elucidation of the molecular mechanism of the breakage-fusion-bridge (BFB) cycle using a CRISPR-dCas9 cellular model. Nucleic Acids Res, 2024.

55. Porro, A., et al., Functional characterization of the TERRA transcriptome at damaged telomeres. Nat Commun, 2014. 5: p. 5379.

56. Nergadze, S.G., et al., CpG-island promoters drive transcription of human telomeres. RNA, 2009. 15(12): p. 2186–94.

57. Arnoult, N., A. Van Beneden, and A. Decottignies, Telomere length regulates TERRA levels through increased trimethylation of telomeric H3K9 and HP1alpha. Nat Struct Mol Biol, 2012. 19(9): p. 948–56.

58. Deng, Z., et al., A role for CTCF and cohesin in subtelomere chromatin organization, TERRA transcription, and telomere end protection. EMBO J, 2012. 31(21): p. 4165–78.

59. Feretzaki, M., P. Renck Nunes, and J. Lingner, Expression and differential regulation of human TERRA at several chromosome ends. RNA, 2019. 25(11): p. 1470–1480.

60. Rivosecchi, J., K. Jurikova, and E. Cusanelli, Telomere-specific regulation of TERRA and its impact on telomere stability. Semin Cell Dev Biol, 2024. 157: p. 3–23.

61. Kyriacou, E. and J. Lingner, TERRA long noncoding RNA: At the interphase of telomere damage, rescue and signaling. Curr Opin Cell Biol, 2024. 91: p. 102437.

62. Silva, B., et al., TERRA transcription destabilizes telomere integrity to initiate break-induced replication in human ALT cells. Nat Commun, 2021. 12(1): p. 3760.

63. Guh, C.Y., et al., XPF activates break-induced telomere synthesis. Nat Commun, 2022. 13(1): p. 5781.

64. Xu, M., et al., TERRA-LSD1 phase separation promotes R-loop formation for telomere maintenance in ALT cancer cells. Nat Commun, 2024. 15(1): p. 2165.

