## Supplemental Materials for "DNA2 and FANCM function in two distinctive pathways in disrupting TERRA R-loops and suppressing replication stress at ALT telomeres"

**Supplemental Information**

**Supplemental Figure Legends**

**Figure S1**. Cell cycle analysis of siRNA transfected U2OS cells.

**Figure S2**. Inhibition of DNA2 using a small molecule inhibitor, d16, in ALT+ cells induces a pronounced replication stress response at their telomeres. U2OS cells were first treated with 10 μM d16 for 48 hours and 72 hours. Cells were then stained with antibodies recognizing TRF2 and pChk1 (A and B), or TRF1 and pRPA (C and D), or TRF1 and γH2AX (E and F). All nuclei were stained with DAPI (Blue). More than two hundred cells were counted for each sample. All error bars are standard deviations obtained from three different experiments. Standard two-sided t test: *$p$<0.05, **$p$<0.01, ***$p$<0.001.

**Figure S3**. Inhibition of DNA2 using a small molecule inhibitor, C5, in ALT+ cells induces a pronounced replication stress response at their telomeres. U2OS cells were first treated with 100 μM C5 for 48 hours and 72 hours. Cells were then stained with antibodies recognizing TRF2 and pChk1 (A and B), or TRF1 and BLM (C and D). All nuclei were stained with DAPI (Blue). More than two hundred cells were counted for each sample. All error bars are standard deviations obtained from three different experiments. Standard two-sided t test: *$p$<0.05, **$p$<0.01, ***$p$<0.001.

**Figure S4**. Depletion of DNA2 in Saos2 cells induces a pronounced replication stress response at their telomeres and an increase of multiple ALT properties. Saos2 cells were first transfected with siLuc or three different siRNA targeting DNA2 (siDNA2-1, siDNA2-2, and siDNA2-3). (**A**) siRNA transfected Saos2 cell lysates were collected and used for immunoblotting (IB). (**B** to **G**) siRNA transfected Saos2 cells were stained with antibodies recognizing TRF1 and pChk1 (B and C), or TRF1 and γH2AX (D and C), or TRF1 and PML (F and G). All nuclei were stained with DAPI (Blue). More than two hundred cells were counted for each sample. All error bars are standard deviations obtained from three different experiments. Standard two-sided t test: *$p$<0.05, **$p$<0.01, ***$p$<0.001.

**Figure S5**. DNA2 deficiency induced DNA damage response at ALT telomeres requires both BRCA1 and BLM. siRNA transfected U2OS cells were stained with antibodies recognizing TRF2 and pChk1 (A and E), or TRF2 and γH2AX (B and F), or TRF2 and pRPA (C and G), or TRF2 and BLM (D). All nuclei were stained with DAPI. More than two hundred cells were counted for each sample. All error bars are standard deviations obtained from three different experiments. Standard two-sided t test: **p*<0.05, ***p*<0.01, ****p*<0.001.

**Figure S6**. DNA2 and FANCM manifest a strong additive genetic interaction in suppressing replication stress response at telomeres in Saos2 cells. siRNA transfected Saos2 cells were stained with antibodies recognizing TRF1 and pChk1 (A), or TRF1 and γH2AX (B), or TRF1 and pRPA (C), or TRF1 and BLM (D). All nuclei were stained with DAPI. More than two hundred cells were counted for each sample. All error bars are standard deviations obtained from three different experiments. Standard two-sided t test: **p*<0.05, ***p*<0.01, ****p*<0.001. (G and H) Saos2 cells were first transfected with different siRNA. DNA was then extracted and used for C-circle assay. "–Φ" indicates samples with no Phi(Φ)29 DNA polymerase added. Image J was used for the quantification of the images. All error bars are standard deviations obtained from three different experiments. Standard two-sided t test: **p*<0.05, ***p*<0.01, ****p*<0.001.

**Figure S7**. All the SMTA-OM molecules of the representative chromosome arm 14q from (A) siLuc and (B) siDNA2-3 transfected U2OS cells.

**Supplemental Tables**:

**Supplemental Table 1**: SMTA-OM analysis of DNA2 and FANCM deficient U2OS and Saos2 cells

**Supplemental Table 2**: SMTA-OM analysis of chromosome arm correlations of DNA2 and FANCM deficient U2OS and Saos2 cells

**Supplemental Table 3**: siRNA list

**Supplemental Table 4**: Antibody list

**Fig S1**

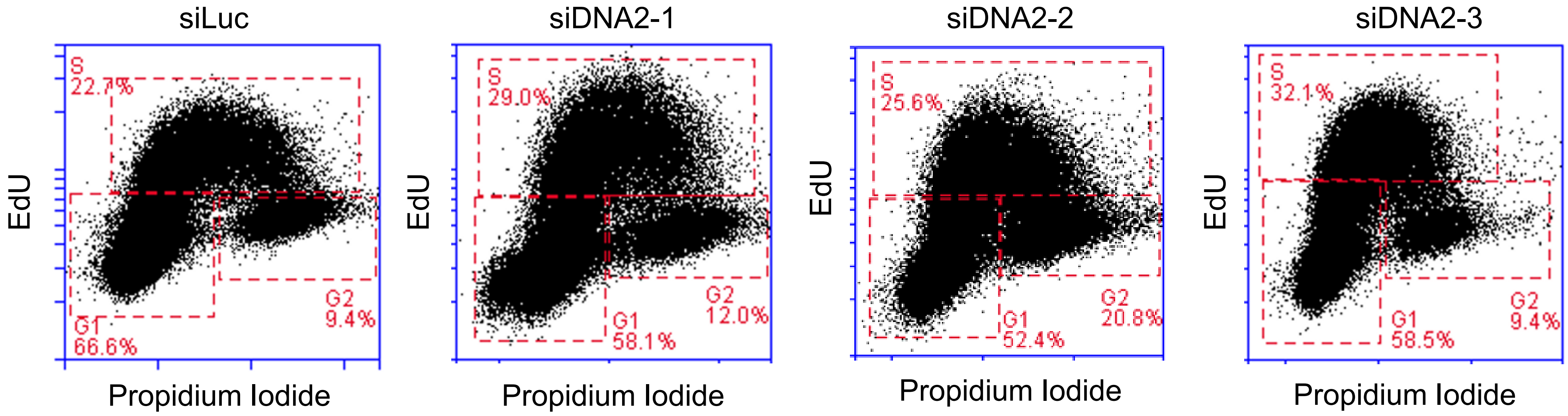


| | **G1** | **S** | **G2** |
|---|---|---|---|
| siLuc | 66.6 | 22.7 | 9.4 |
| siDNA2-1 | 58.1 | 29 | 12 |
| siDNA2-2 | 52.4 | 25.6 | 20.8 |
| siDNA2-3 | 58.5 | 32.1 | 9.4 |

Fig S2

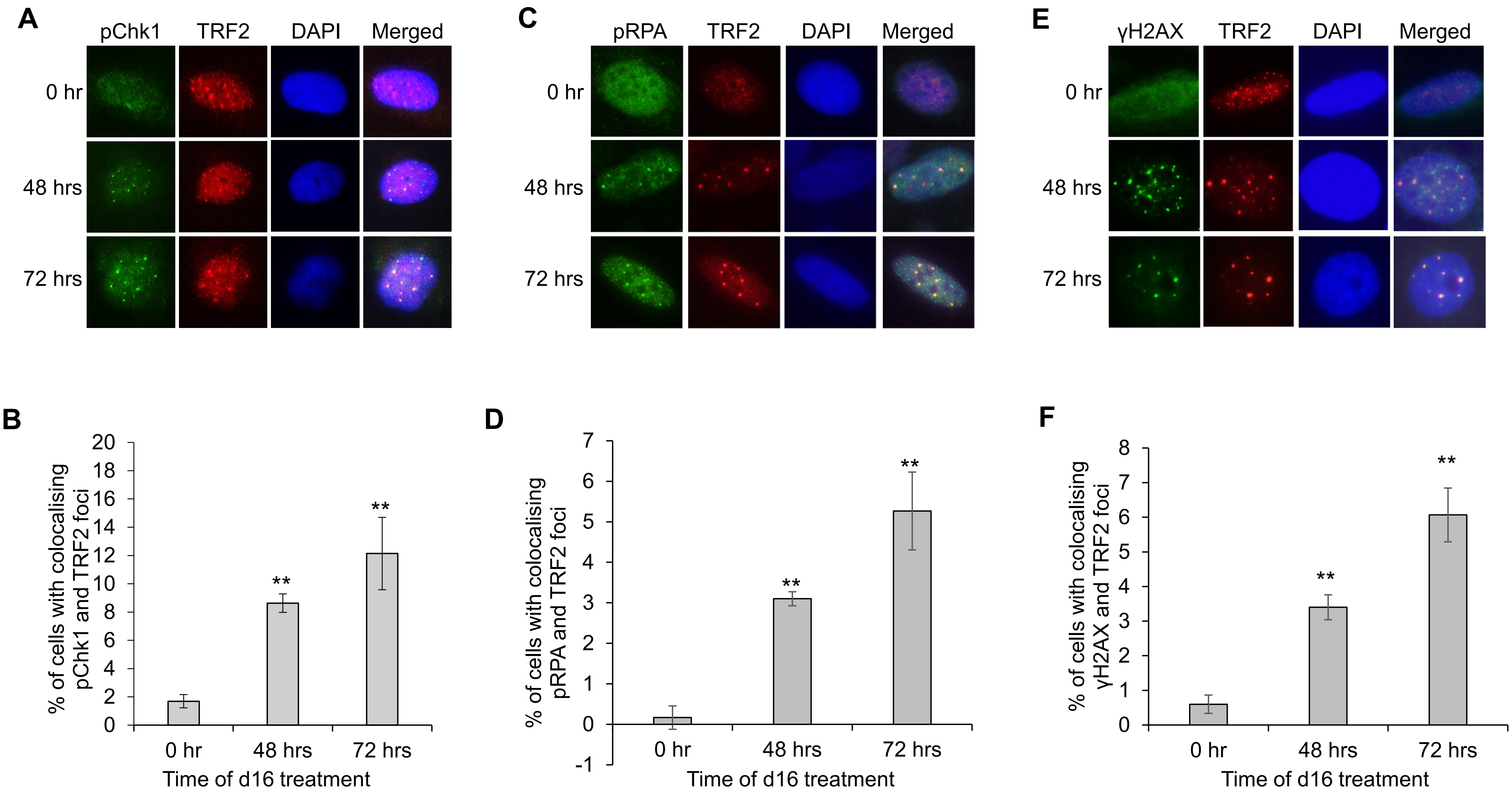
A
pChk1
TRF2
DAPI
Merged
0 hr
48 hrs
72 hrs
C
pRPA
TRF2
DAPI
Merged
0 hr
48 hrs
72 hrs
E
γH2AX
TRF2
DAPI
Merged
0 hr
48 hrs
72 hrs
B
% of cells with colocalising pChk1 and TRF2 foci
20
18
16
14
12
10
8
6
4
2
0
**
**
0 hr
48 hrs
72 hrs
Time of d16 treatment
D
% of cells with colocalising pRPA and TRF2 foci
7
6
5
4
3
2
1
0
-1
**
**
0 hr
48 hrs
72 hrs
Time of d16 treatment
F
% of cells with colocalising γH2AX and TRF2 foci
8
7
6
5
4
3
2
1
0
**
**
0 hr
48 hrs
72 hrs
Time of d16 treatment

**Fig S3**

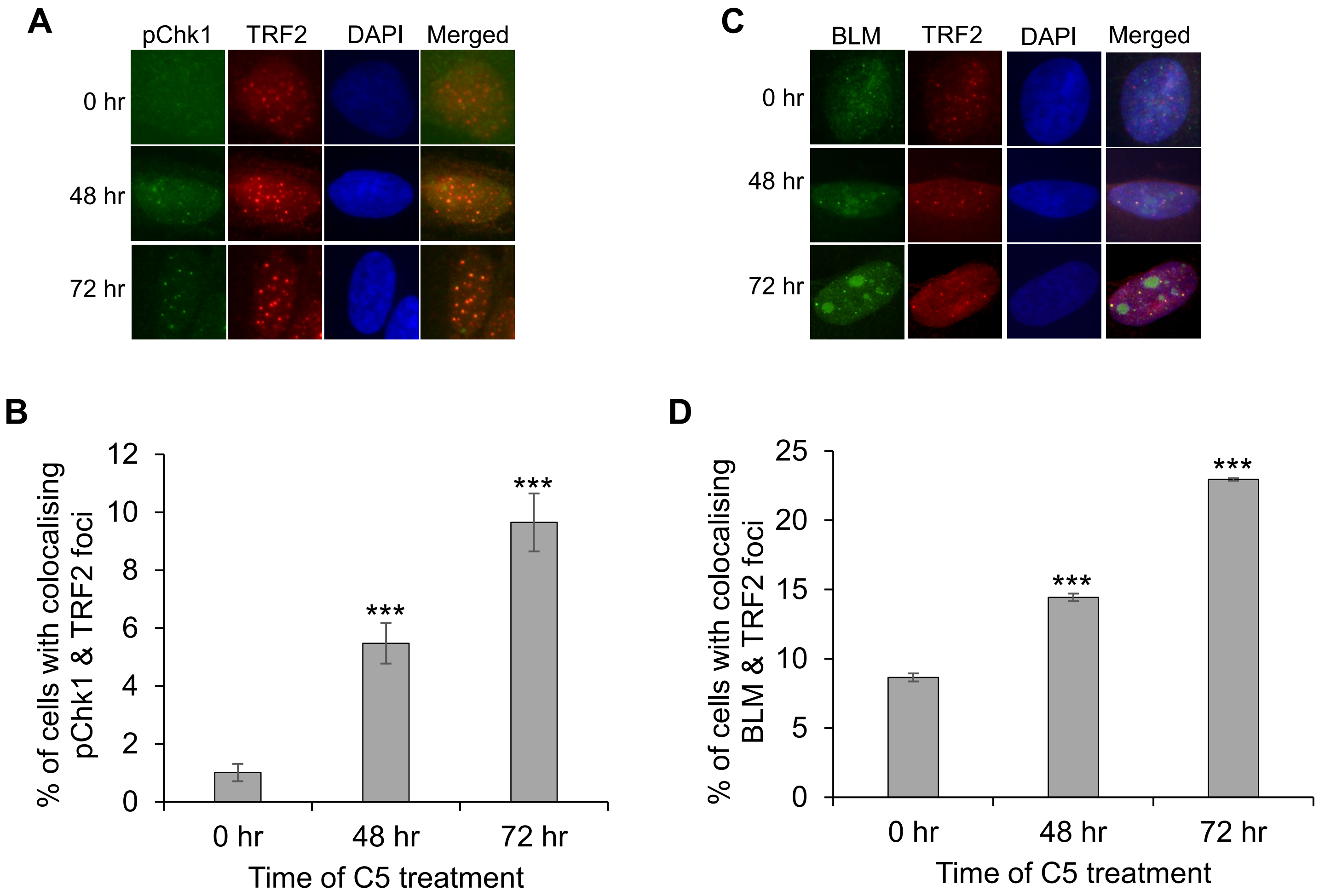
A
pChk1
TRF2
DAPI
Merged
0 hr
48 hr
72 hr
C
BLM
TRF2
DAPI
Merged
0 hr
48 hr
72 hr
B
% of cells with colocalising pChk1 & TRF2 foci
0
2
4
6
8
10
12
***
***
0 hr
48 hr
72 hr
Time of C5 treatment
D
% of cells with colocalising BLM & TRF2 foci
0
5
10
15
20
25
***
***
0 hr
48 hr
72 hr
Time of C5 treatment

Fig S4

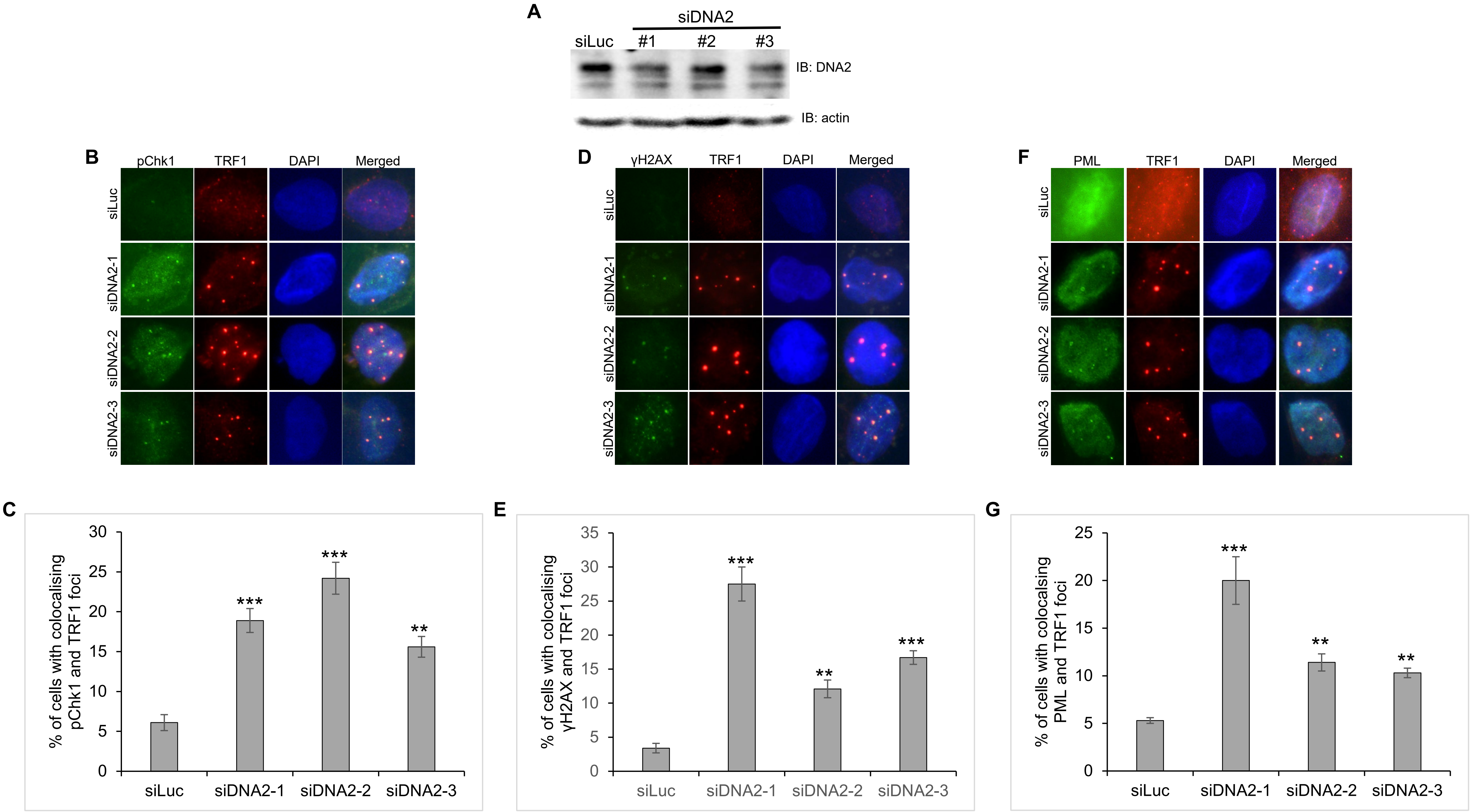
A
siDNA2
siLuc
#1
#2
#3
IB: DNA2
IB: actin
B
pChk1
TRF1
DAPI
Merged
siLuc
siDNA2-1
siDNA2-2
siDNA2-3
D
γH2AX
TRF1
DAPI
Merged
siLuc
siDNA2-1
siDNA2-2
siDNA2-3
F
PML
TRF1
DAPI
Merged
siLuc
siDNA2-1
siDNA2-2
siDNA2-3
C
% of cells with colocalising pChk1 and TRF1 foci
0
5
10
15
20
25
30
***
***
**
siLuc
siDNA2-1
siDNA2-2
siDNA2-3
E
% of cells with colocalising γH2AX and TRF1 foci
0
5
10
15
20
25
30
35
***
**
***
siLuc
siDNA2-1
siDNA2-2
siDNA2-3
G
% of cells with colocalsing PML and TRF1 foci
0
5
10
15
20
25
***
**
**
siLuc
siDNA2-1
siDNA2-2
siDNA2-3

**Fig S5**

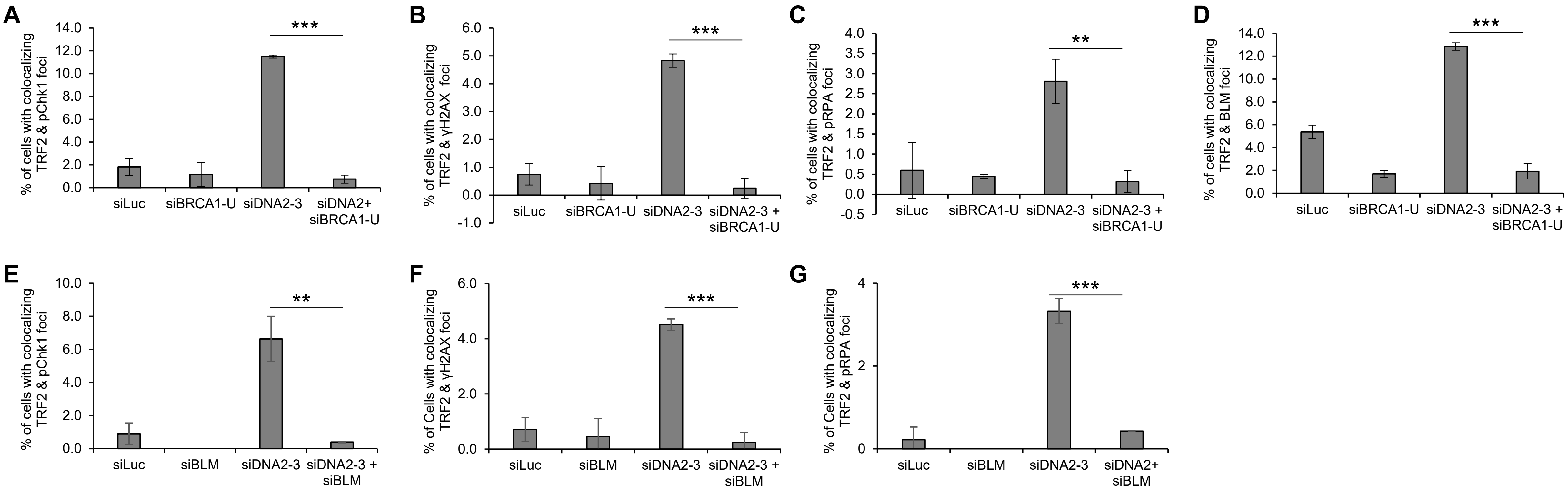
A
% of cells with colocalizing TRF2 & pChk1 foci
14.0
12.0
10.0
8.0
6.0
4.0
2.0
0.0
***
siLuc
siBRCA1-U
siDNA2-3
siDNA2+ siBRCA1-U
B
% of cells with colocalizing TRF2 & γH2AX foci
6.0
5.0
4.0
3.0
2.0
1.0
0.0
-1.0
***
siLuc
siBRCA1-U
siDNA2-3
siDNA2-3 + siBRCA1-U
C
% of cells with colocalizing TRF2 & pRPA foci
4.0
3.5
3.0
2.5
2.0
1.5
1.0
0.5
0.0
-0.5
**
siLuc
siBRCA1-U
siDNA2-3
siDNA2-3 + siBRCA1-U
D
% of cells with colocalizing TRF2 & BLM foci
14.0
12.0
10.0
8.0
6.0
4.0
2.0
0.0
***
siLuc
siBRCA1-U
siDNA2-3
siDNA2-3 + siBRCA1-U
E
% of cells with colocalizing TRF2 & pChk1 foci
10.0
8.0
6.0
4.0
2.0
0.0
**
siLuc
siBLM
siDNA2-3
siDNA2-3 + siBLM
F
% of Cells with colocalizing TRF2 & γH2AX foci
6.0
4.0
2.0
0.0
***
siLuc
siBLM
siDNA2-3
siDNA2-3 + siBLM
G
% of Cells with colocalizing TRF2 & pRPA foci
4
2
0
***
siLuc
siBLM
siDNA2-3
siDNA2+ siBLM

Fig S6

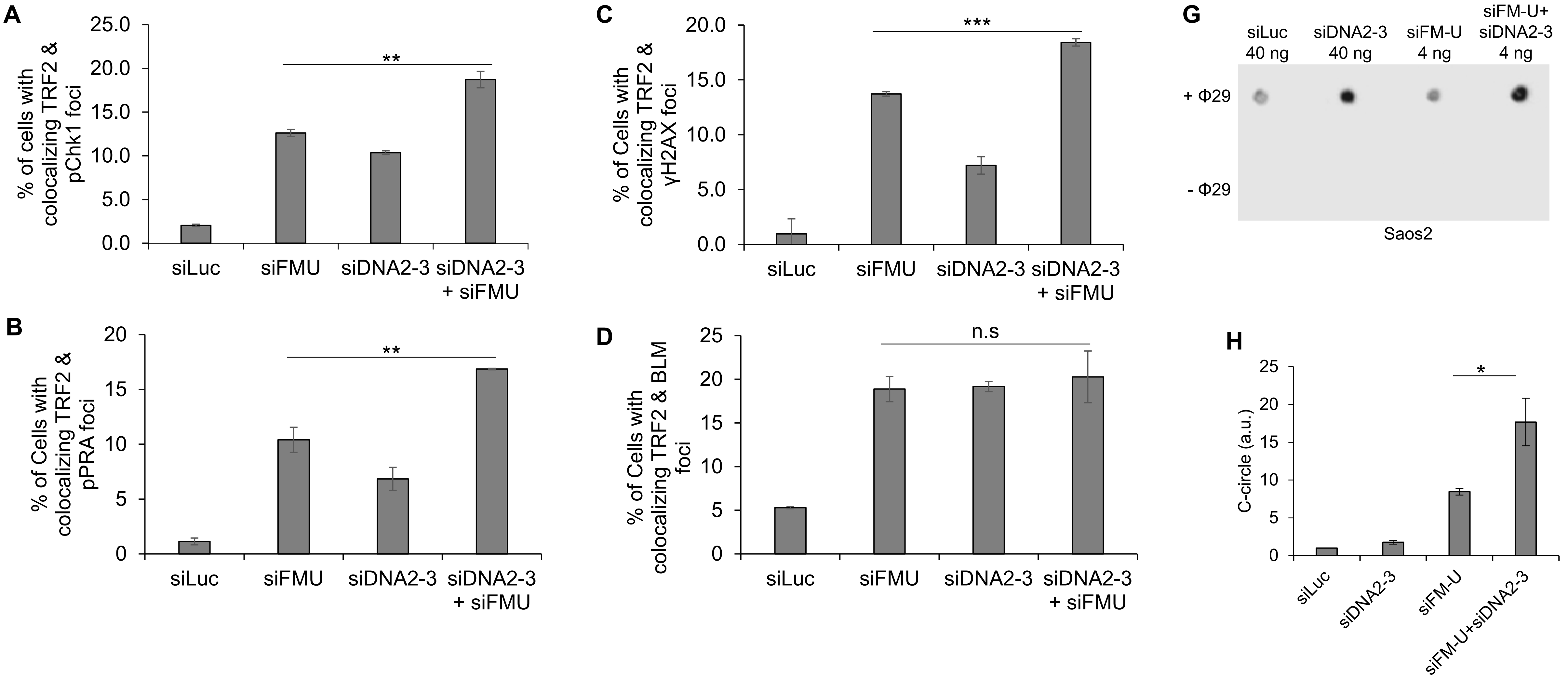
A
% of cells with colocalizing TRF2 & pChk1 foci
25.0
20.0
15.0
10.0
5.0
0.0
**
siLuc
siFMU
siDNA2-3
siDNA2-3 + siFMU
B
% of Cells with colocalizing TRF2 & pPRA foci
20
15
10
5
0
**
siLuc
siFMU
siDNA2-3
siDNA2-3 + siFMU
C
% of Cells with colocalizing TRF2 & γH2AX foci
20.0
15.0
10.0
5.0
0.0
***
siLuc
siFMU
siDNA2-3
siDNA2-3 + siFMU
D
% of Cells with colocalizing TRF2 & BLM foci
25
20
15
10
5
0
n.s
siLuc
siFMU
siDNA2-3
siDNA2-3 + siFMU
G
siLuc 40 ng
siDNA2-3 40 ng
siFM-U 4 ng
siFM-U+ siDNA2-3 4 ng
+ Φ29
- Φ29
Saos2
H
C-circle (a.u.)
25
20
15
10
5
0
*
siLuc
siDNA2-3
siFM-U
siFM-U+siDNA2-3

Fig S7

A

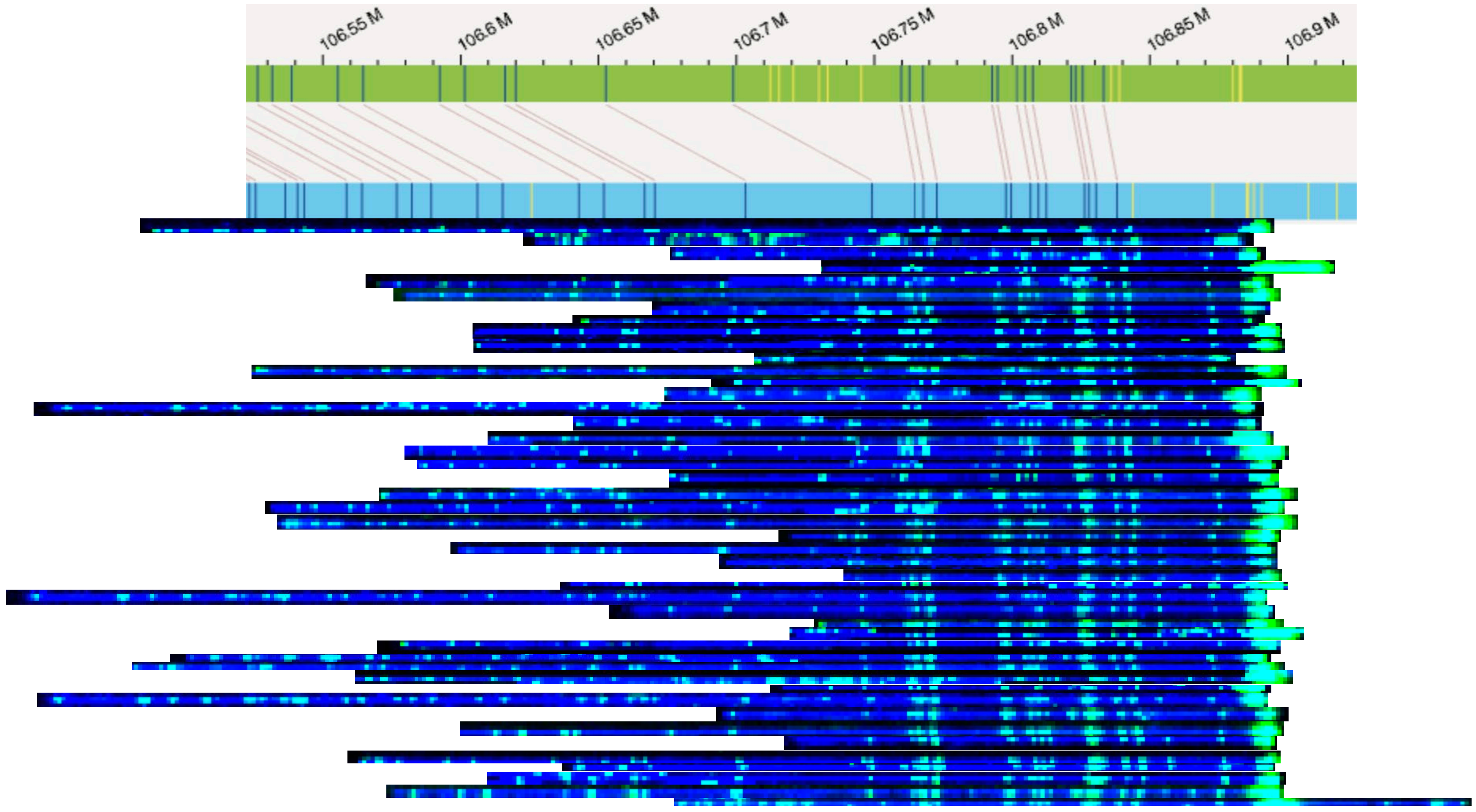
106.55 M
106.6 M
106.65 M
106.7 M
106.75 M
106.8 M
106.85 M
106.9 M

B

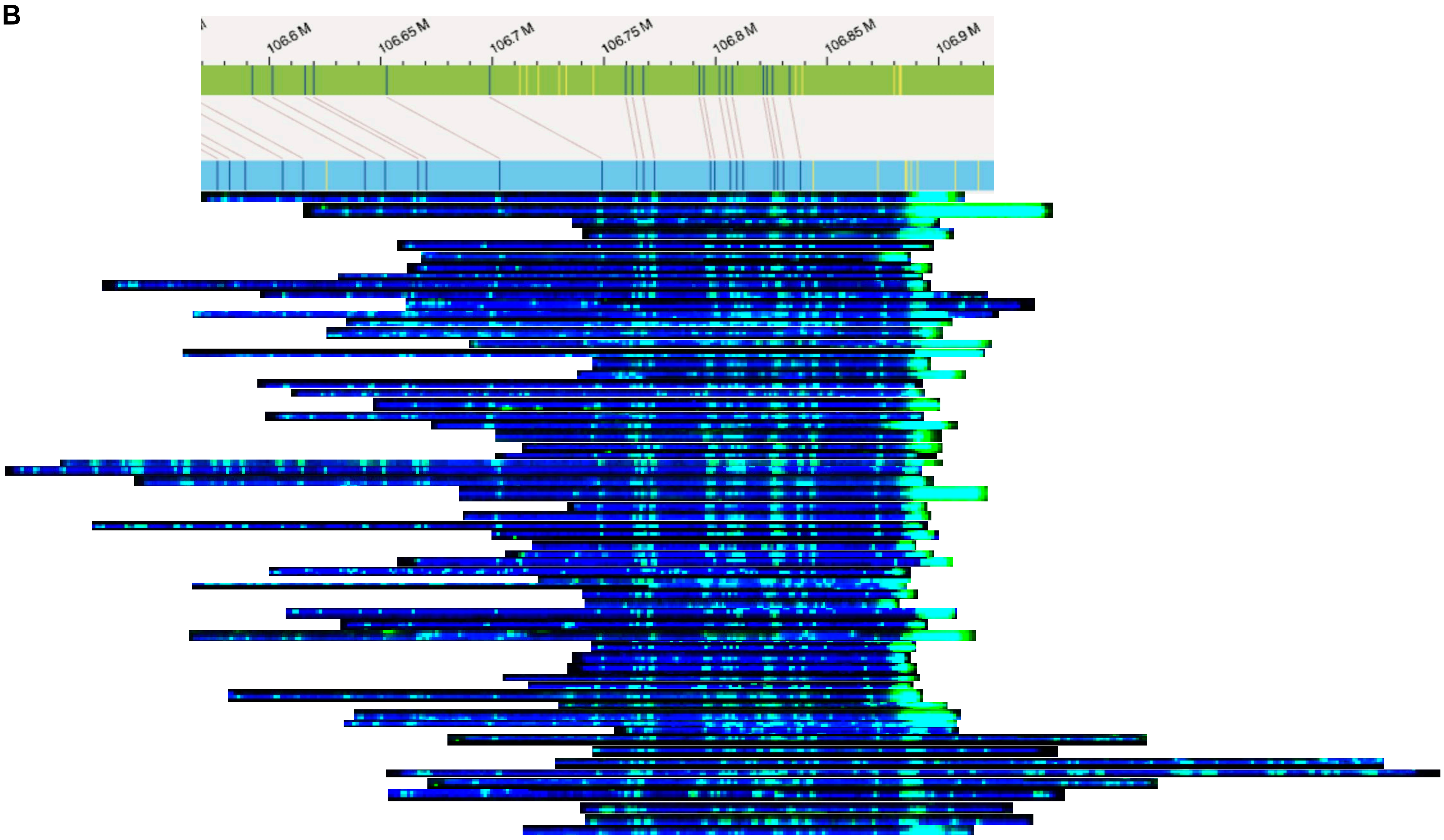
106.6 M
106.65 M
106.7 M
106.75 M
106.8 M
106.85 M
106.9 M
